# Transient ion-mediated interactions regulate subunit rotation in a eukaryotic ribosome

**DOI:** 10.1101/2025.08.09.669508

**Authors:** George Wanes, Udayan Mohanty, Paul Whitford

## Abstract

While it is known that ions are required for folding of RNA, little is known about how transient/probabilistic ionic interactions facilitate biologically-relevant conformational rearrangements. To address this, we developed a theoretical model that employs all-atom resolution, with a simplified representation of biomolecular energetics, explicit electrostatics and ions (K^+^, Cl^−^, Mg^2+^). For well-studied RNA systems (58-mer and Ade riboswitch), the model accurately describes the concentration-dependent ionic environment, including (bidentate) chelated and hydrated (diffuse/outer-shell) ions. With this foundation, we applied the model to simulate the yeast ribosome and quantified the ion-dependent energy landscape of intersubunit rotation. These calculations show how the energetics of rotation responds to millimolar changes in [MgCl_2_], which shift the distribution between rotation states and alter the kinetics by more than an order of magnitude. We find that this response to the ionic concentration correlates with formation and breakage of ion-mediated interactions (inner-shell and outer-shell) between the ribosomal subunits. This analysis provides a physical basis for understanding how transient ion-mediated interactions can regulate a large-scale biological process.

## Introduction

Cations are indispensable for a wide range of cellular processes.^1,2^ K^+^ and Mg^2+^ are the most abundant cations in the cell,^3^ and they ensure proper assembly, folding and dynamics of RNA,^4–8^ DNA^9,10^ and ribonucleoprotein (RNP)^11^ complexes. They contribute to catalytic steps,^12,13^ facilitate folding of RNA by screening repulsive electrostatic interactions and stabilize negatively-charged backbone interactions.^8^ Numerous examples of these effects can be found in the ribosome, a complex RNP assembly that requires precise ionic conditions for folding^14,15^ and translation.^16,17^ While high-resolution structures obtained with X-ray crystallography^14,18,19^ and cryo-EM^20–22^ can identify highly-stable (i.e. “structural”) ions, determining the precise ways that transient ionic interactions enable large-scale conformational rearrangements has remained elusive.

Probing the interplay between ion and RNA dynamics is a challenge, in part, because the interactions occur over a wide range of time and length scales. Mg-RNA interactions may be described in terms of three general modes (Fig. 1a): inner-shell, outer-shell and bulk. Inner-shell/chelated interactions are associated with the shortest length scales (~ 2Å), and their formation requires release of a water molecule, which is energetically unfavorable.^6^ Once formed, these strong interactions are long-lived (ms-s timescales).^23,24^ Inner-shell interactions may be further categorized as unidentate (i.e. chelated with a single atom; Fig. 1a), or multidentate (bridging atoms in multiple residues; Fig. 1b), where the latter are often thought of as “permanent” components of a molecular structure.^14^ In contrast, outer-shell/diffuse ions remain fully hydrated and are associated with longer-range (~ 4 − 5Å) RNA interactions that are estimated to remain formed for shorter time scales.^7,25^ Despite the fact that their interactions are weaker, high concentrations of diffuse ions can form near RNA, allowing them to provide a significant contribution to molecular energetics.^25,26^ Farther from RNA (*>* 8Å), bulk ions can contribute to long-range electrostatic screening^7^ by organizing a dispersed counter-ionic atmosphere.

**Figure 1:**
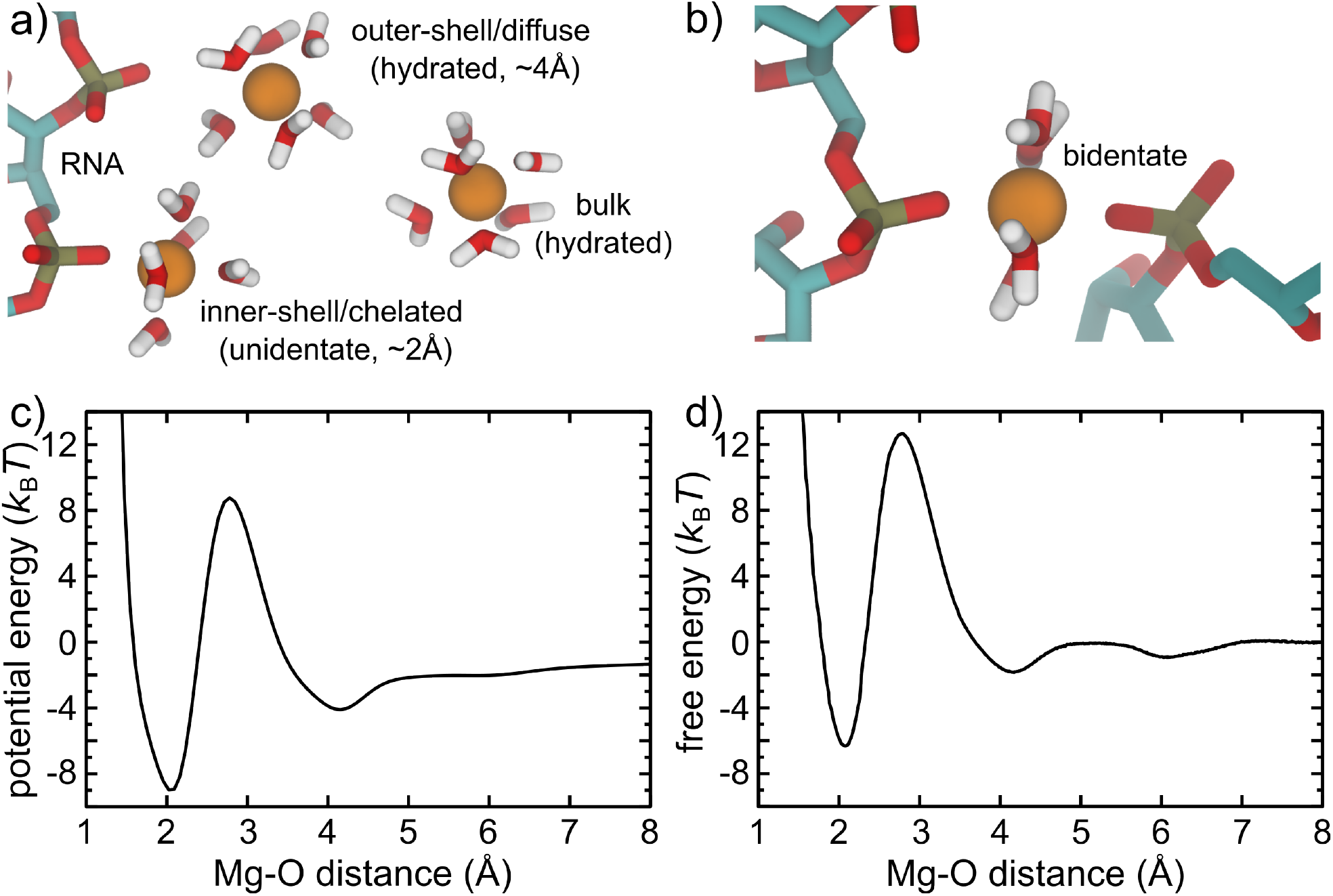
Describing the energetics of Mg^2+^-RNA interactions. Mg^2+^ ions contribute to the functional dynamics in RNA at multiple length and time scales. a) While remaining fully hydrated, Mg^2+^ ions can transiently (ns-*µ*s) interact with RNA through long-range electrostatics in bulk (~ 10Å), or condense around RNA to form the so-called “outer-shell” (~ 4Å). Longer-lived (ms) short-range (~2Å) inner-shell/chelated interactions (unidentate) can be formed with highly-charged atoms (e.g. non-bridging phosphate oxygens). b) A single Mg^2+^ ion can also form multiple inner-shell interactions (bidentate/multidentate) between RNA residues that are distant in sequence. These extremely stable states often thought of as “permanent” components of the molecular architecture. c) The current study introduces a model (SMOG+inner-ion) that employs an effective potential for Mg^2+^-O interactions that quantitatively describes these four modes of Mg^2+^-RNA interactions. d) The free-energy as a function of Mg^2+^-O distance, calculated with this model for a reference compound (see methods). Consistent with available structural data, there is a deep minimum at 2.07 Å, corresponding to the stable chelated state. The barrier to escape the inner-shell/chelated state is ~ 19 k_*B*_T, which is consistent with experimental dissociation timescales (milliseconds). The corresponding binding affinity is −1.2 k_*B*_T, also consistent with experimental estimates.

There has been progress identifying how and when different modes of Mg^2+^ interactions are utilized in biological systems, though significant obstacles remain. Structural methods often provide ambiguous Mg^2+^ assignments, since water molecules and Na^+^ ions have similar electron densities.^27^ This limitation is exacerbated when analyzing lower-resolution structures, where the hydration shell around each ion may not be visualized. To help address this, stereochemical guidelines^28–30^ and machine learning techniques^31,32^ have been developed that can identify signatures of ions in cryo-EM reconstructions. In recent studies, these guidelines were used to reassess a cryo-EM structure of a bacterial ribosome,^33^ which provided a more systematic identification of ion binding sites within the core of the ribosomal RNA. Although these represent significant advances, the methods rely on experimentally obtaining sufficient density for each ion. which is not always possible in more dynamic structural elements.^14^

Energetic models can expand our understanding of ions by describing the probabilistic nature of the interactions. At the highest level of resolution are Quantum Mechanical (QM) calculations that account for the electronic environment.^34,35^ However, a practical challenge when studying large systems is that these methods can only be applied to small subsets of atoms.^35–37^ In addition, there can be thousands of possible ion binding sites in large RNPs, and even a single QM calculation can be extremely computationally demanding.^34,38^ To complement QM methods, explicit-solvent force fields have been developed^39–41^ that provide high spatial resolution, where the accessible timescales are typically limited to microseconds. In addition, the predicted kinetics of ion-RNA interactions are often model-specific,^40–44^ which introduce additional uncertainties when interpreting dynamics. At a coarse-grained level, implicit-solvent models that apply Debye-Hückel (DH) theory^45–47^ or the reference interaction site model^48–50^ have enabled simulations of larger systems at a reduced computational cost. While these approaches often neglect ion-ion correlations, there are strategies for introducing explicit divalent ions with implicit monovalent ion treatments.^50–52^ Unfortunately, coarse-graining of RNA structure can introduce artificial steric effects that can strongly influence conformational motions in RNPs.^53^ Simplified models that apply all-atom representations provide more complete steric descriptions at lower computational costs, and they have enabled the simulation of large-scale rearrangements^54^ and folding^55^ with ion-free models. Building on this, we previously developed a model with higher molecular resolution, as well as monovalent and divalent ions, though it only accounted for ions in the outer-shell/diffuse and bulk states.^26^

To study how transient inner-shell and outer-shell ions contribute to the energetics of large-scale biomolecular rearrangements, we introduce a model that applies an all-atom representation of the biomolecule, along with explicit ions (Mg^2+^, K^+^, Cl^−^). In this model, intramolecular interactions are provided by a structure-based force field,^56^ while ionic interactions are described by direct Coulomb potentials and effective potentials that account for ion-hydration effects. To complement experimental techniques that can resolve bound ions in highly stable conformations, this model can be used to probe the dynamics of ion-mediated interactions during large-scale conformational transitions. Below, we first show that the force field accurately describes the ionic environment around model compounds and well-studied RNA molecules. We then characterize the energetics and kinetics of subunit rotation in the yeast ribosome. The simulations uncover an ion-concentration-dependent energy landscape, where site-specific ion-mediated interactions dynamically reorganize during this large-scale collective rearrangement.

## Methods

### Structure-based “SMOG” model with electrostatics and explicit inner and outer-shell ions

In the presented force field (called the SMOG+inner-ion model), all non-hydrogen atoms, Mg^2+^, K^+^ and Cl^−^ are explicitly represented. Biomolecular interactions are defined to stabilize an assigned structure (i.e. a structure-based “SMOG”^56,57^ model). To include Coulomb effects in the biomolecule/s, partial charges are assigned to all atoms. Ions interact through Coulomb potentials that are augmented by terms that account for solvation effects. Parameters are the same as those in our previous diffuse-only ion model (“SMOG-ion”),^26^ except for modifications that were introduced to describe inner-shell Mg^2+^ and K^+^ interactions with RNA. For completeness, the full force field is described here and parameterization steps are given in the next section.

The potential energy may be expressed as:

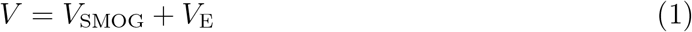

The first term represents the all-atom structure-based model^56^ and the functional form is given by:

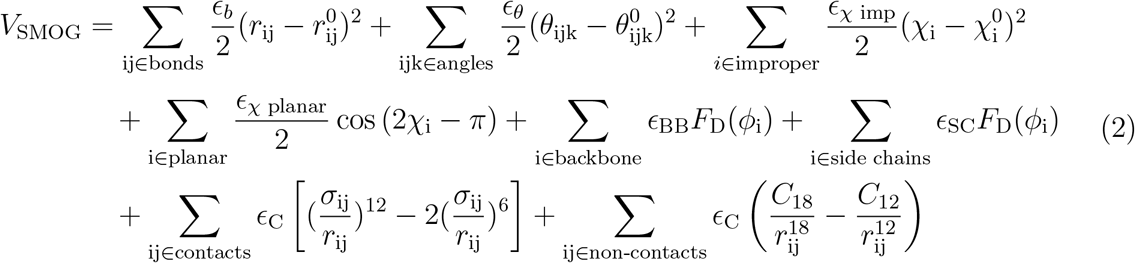

where

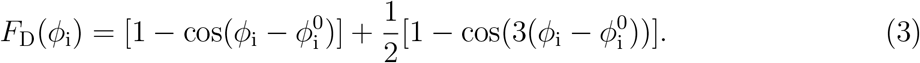

*V* _SMOG_ describes the intermolecular and intramolecular interactions of the biomolecule/s, where a preassigned configuration is defined as the global potential energy minimum. Bonded parameters 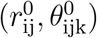 are given the values found in the Amber99sb-ildn force field,^58^ as described previously.^59^ Dihedral 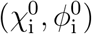 and contact (*σ*_ij_) parameters are assigned the values found in the experimental configuration, unless otherwise specified. The Shadow Contact Map algorithm^60^ is used to define the contacts, with the default 6 Å cutoff and 1Å shadowing radius. The 12-18 term represents the excluded volume for atom pairs that are not in contact in the experimental structure. As described previously,^26^ the C_18_ and C_12_ parameters are defined to mimic the repulsive component of 6-12 interactions in Amber99sb-ildn force field.^58^

The second term in Eq. 1 describes electrostatic and solvation effects, and it is given by:

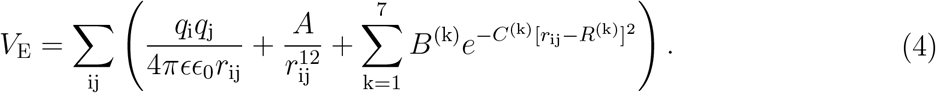

*q*_i_ and *q*_j_ are the atomic charges, which are defined based on the Amber99sb-ildn force field.^58^ Because the SMOG+inner-ion model does not include hydrogen atoms, the partial charge of each hydrogen atom is added to the corresponding non-hydrogen atom. *E* is the dielectric constant of water (80), and *E*_0_ is the permittivity of free space. The parameters for the excluded volume terms (*A*) are defined separately for each interaction type (e.g., Mg-Mg, Mg-Cl, Mg-O, etc.). The final term is the ionic desolvation term, which is given by a sum of Gaussians (Fig. 1c) that describe both the inner and outer ionic shells.

Since SMOG models do not include the full energetic roughness of a system, we use diffusion coefficients to estimate the effective simulated time unit.^61^ The effective time unit (*τ*^eff^) is estimated by equating the diffusion coefficients calculated with the SMOG model (in reduced units) and an experimental values and/or the value predicted by an explicit-solvent model (e.g. CHARMM,^62^ AMBER^63^). The rationale for this approach is that, while the large-scale properties of the energy landscape can be described by a SMOG model, these models lack the full degree of non-specific roughness that reduces the rate of diffusion.^64,65^ Accordingly, by comparing diffusion coefficients with experiments or explicit-solvent models, one is able to estimate the speedup in a SMOG model and the effective simulated time. With the SMOG+inner-ion model and simulation parameters, the self-diffusion coefficient of Mg^2+^ is ~ 47 Å^2^/*τ*^eff^ (Fig. S2). Equating this with the experimental value of *D*_0_ = 0.71 *×* 10^−5^ cm^2^/s^66^ indicates that the effective simulated time unit, *τ*^eff^, corresponds to approximately 66 ps. This is within the range estimated from simulations with an electrostatics-free SMOG model and an explicit-solvent model for tRNA within the accommodation corridor of the ribosome (50 ps - 1 ns).^67^ Accordingly, we use an effective time unit of 66 ps when reporting simulated times with the SMOG+inner-ion model.

### Parameterizing the inner-shell ion potential

To parameterize the effective potentials that describe inner-shell ion-RNA interactions, the following steps were performed. All analysis presented in the Results section describes predictions that were produced after all refinement steps were completed.

1. The ion excluded volume *{A}*, outer-shell Mg^2+^ parameters and inner-shell K-O parameters were refined against explicit-solvent simulations using the iterative process method described by Savelyev and Papoian,^68^ as applied when developing the previous SMOG-ion model.^26^ In the SMOG-ion model, inner-shell K-O interactions were excluded after refinement. In the SMOG+inner-ion model, we use the refined parameters determined previously, without removal of the K-O inner-shell potential. For Mg-P interactions, *A* was given a sufficiently large value that prevents the formation of bidentate interactions with a single phosphate group (Fig. S1).
2. Inner-shell Mg-O parameters were manually adjusted in order to accurately capture experimental rates and configurations. The inner-shell interactions are described by the first two Gaussians terms in equation 4. The stability of the Mg-O inner-shell interaction is defined by *B*^(1)^ and the desolvation barrier height is set by *B*^(2)^. The positions (*R*^(1)^) and width (*C*^(1)^) of the inner-shell were manually adjusted in order for the free-energy to have a minimum at 2.07Å (Fig. 1d), consistent with available structural data^69 70^.^71^ *B*^(1)^ was calibrated to predict the binding constant of Mg^2+^ with O atoms, as interpolated by Sigel and Sigel using potentiometric pH titration experiments.^72^ The height of the desolvation barrier (*B*^(2)^) was assigned a value that produced a dissociation rate for Mg^2+^ binding of ≈ 10^3^−10^4^*s*^−1^, as implicated by experimental measurement using P-NMR,^24,73^^25^Mg NMR,^23^ variable-temperature NMR^74^ and temperature-jump spectroscopy^75^ for a range of RNA and DNA systems.
3. The depth of the outer-shell interaction (*B*^(3)^) was adjusted to ensure a proper balance between inner-shell and outer-shell energetics. After an initial parameter sweep, we used a reweighting strategy^26^ to estimate a value of *B*^(3)^ that would produce the experimental value of the preferential interaction coefficient (Γ_2+_) of Mg^2+^ ions for the 58-mer at [MgCl_2_] = 1mM and [KCl] = 150mM. We then repeated the simulations with this value of *B*^(3)^, in order to confirm that the model reproduces the experimental value of Γ_2+_ under the reference ionic conditions.

### Multi-basin structure-based model for the ribosome

To study the energetics of subunit rotation in the yeast ribosome, a multi-basin SMOG model was used. Following the protocol used previously,^76^ the minimum for each dihedral angle 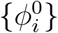 was set to the mean periodic value between the unrotated (RCSB ID: 3J78) and rotated (RCSB ID: 3J77) conformations,^77^ which avoids bias towards either state. Similarly, for the contacts that are common to both structures, we defined *σ*_i,j_ (the distance at which the 6-12 term is minimal) such that the two conformations are isoenergetic (energy of −*ϵ*_iso_). A contact is defined as “common” if *ϵ*_iso_*/E*_*C*_ *>* 1*/*2 (i.e. small difference in distance in the two structures). For contacts that are unique to either conformation, *σ*_i,j_ was assigned the distance from the structure that contains the contact. Contacts that are unique to the rotated conformation were assigned an energetic weight of 0.16*ϵ*, while those that are unique to the unrotated conformation were given a weight of 0.19*ϵ. ϵ* is the reduced energy scale and it is equal to 2 k_*B*_T. These values were chosen so that the rotated and unrotated states correspond to minima in the free-energy landscape, which allows us to probe the extent to which changes in ionic conditions impact the energetics of rotation.

### Binding affinity calculation

We calculated the binding affinity at standard conditions using the following equation:^78^

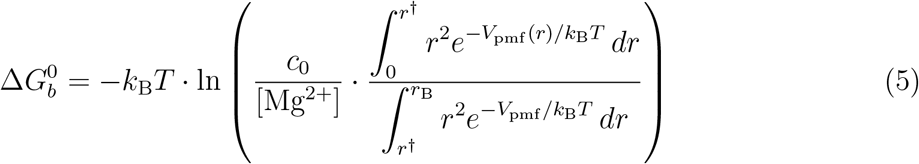

where *r* is the Mg-O distance, *V*_pmf_ is the potential of mean force (free energy; Fig. 1d), *c*_0_ = 1M is the standard reference concentration, *r*^*†*^ is the position of the barrier (2.7 Å) and *r*_B_ is the radius of a sphere with a volume of the simulated box. Simulations with [Mg^2+^] = 3.4 mM were used to calculate the free energy.

## Results

This study describes the physical relationship between short-scale ion-RNA interactions and large-scale dynamics in biomolecular assemblies. We present a model (SMOG+inner-ion) that employs atomistic resolution for each biomolecule and explicit ions (Mg^2+^, K^+^, Cl^−^), where both inner-shell (chelated) and outer-shell (diffuse) interactions are described. While the biomolecular energetics is primarily defined based on knowledge of a specific structure (i.e. Structure-based model^56^), ions interact through Coulomb potentials and effective potentials that account for ionic hydration (Eq. 4). With this model, we first performed a range of benchmark simulations for well-studied systems, which verified that the short-scale geometrical properties, kinetics and energetics of ion-RNA interactions are properly described. We then applied the model to study the dynamics of subunit rotation in the yeast ribosome. These simulations reveal how a combination of inner-shell and outer-shell interactions contribute to the energetics and kinetics of this collective structural rearrangement.

### Modeling inner-shell geometry and energetics of Mg^2+^

To calibrate the model, we assessed the energetic and structural properties of short-scale chelated/inner-shell ion-RNA interactions. We focused on the Mg-O parameters, since Mg^2+^ predominantly interacts with RNA through coordination to oxygen atoms.^28,30,33,79^ As described below, the energetic weights of inner-shell and outer-shell interactions, as well as the height of the intervening barrier (Fig. 1c; Eq. 4) were defined such that the predicted structural and dynamical properties of inner-shell ions are consistent with a range of experimental measurements and ab-initio calculations.

The model was parameterized to accurately describe the structural properties of Mg^2+^ interactions with phosphate-containing model compounds. First, we considered interactions of Mg^2+^ with a single phosphate group. For this, the depth and position of the innershell potential were defined such that the free energy as a function of Mg-O distance has a minimum at 2.07 Å (Fig. 1d). This distance was chosen in order to match the values found in high-resolution structures of fully coordinated (*i*.*e*. hexacoordinated) Mg^2+^ ions. Specifically, for high-resolution entries (R-factor less than 0.05) in the Cambridge Structural Database (CSD),^69,70^ the average Mg-O distance is 2.07 *±* 0.04 Å (ave. *±* s.d.), which is consistent with ab-initio calculations^71,80^ of Mg^2+^ interacting with di-methyl phosphate (2.0 - 2.15 Å). In addition, our model strongly disfavors the formation of intraphosphate bidentate interactions (Fig. S1), consistent with ab-initio calculations^81,82^ and the absence of these interactions in high-resolution structures.^83^ See SI for complete details.

The model was also calibrated to balance the energetics of inner-shell and outer-shell interactions. The weights of the inner- and outer-shell interactions (Fig. 1c) were parameterized to capture an experimentally-measured value of the preferential interaction coefficient Γ_2+_, as well as the binding energy associated with inner-shell/chelated interactions. Γ_2+_ is a common benchmark measure that is used in theoretical studies,^26,50,84^ since it describes the number of divalent ions that are condensed around the RNA, and it is directly related to the energetics of Mg-RNA association.^85^ With the model, the binding free energy of an inner-shell Mg^2+^ with an isolated phosphate group is −1.2 *k*_B_*T* (see Methods), which is similar to the value of −1.04 *k*_B_*T* that has been estimated for Mg^2+^-phosphate using potentiometric pH titration experiments.^72^ Since Γ_2+_ is sensitive to both the inner- and outer-shells, we tuned the outer-shell energetics in the model, in order to capture Γ_2+_ for a model system. For this step, we used a 58-mer RNA with [MgCl_2_] = 1 mM and [KCl] = 150 mM, and the calculated of Γ_2+_ was 10.4, as reported experimentally.^86^

To calibrate the potential energy barrier that separates the inner and outer-shell (Fig. 1c), we considered the dissociation kinetics of Mg^2+^ from a phosphate oxygen. With the model parameters, the dissociation rate was 2.5 *×* 10^3^s^−1^ (See Fig. S3 and SI Results), which is within the experimentally measured range of ≈ 10^3^ − 10^4^s^−1^.^23,24,73–75^ The resulting free-energy barrier for dissociation is ~19 *k*_B_*T* (Fig. 1d), again similar to estimates from explicit-solvent models.^41,87^ Combined with the above comparisons, these benchmarks ensure that the model provides experimentally-consistent distributions and kinetics of inner-shell and outer-shell ion-RNA interactions.

### Balancing monovalent and divalent ion-RNA interactions

After calibrating the model for small model compounds, we tested whether it accurately describes the ionic environment and predicts the binding sites of intermediate-sized (~50 residue) RNA molecules. We focused on two well-studied systems: the U1061A variant of a 58-mer rRNA fragment^88^ (Fig. 2a) and an adenine riboswitch^89^ (Fig. 2d). We first compared the predicted values of the preferential interaction coefficient Γ_2+_ over a range of Mg^2+^ concentrations (~ 0.1 - 1mM) and multiple values of [KCl] (50 and 150 mM). Since Γ_2+_ describes the number of Mg^2+^ ions that accumulate around an RNA molecule (above the bulk value),^85^ it provides a global measure of the local ionic environment. As described below, the model accurately provides site-specific, as well as concentration-dependent, descriptions of these intermediate-scale RNA systems.

**Figure 2:**
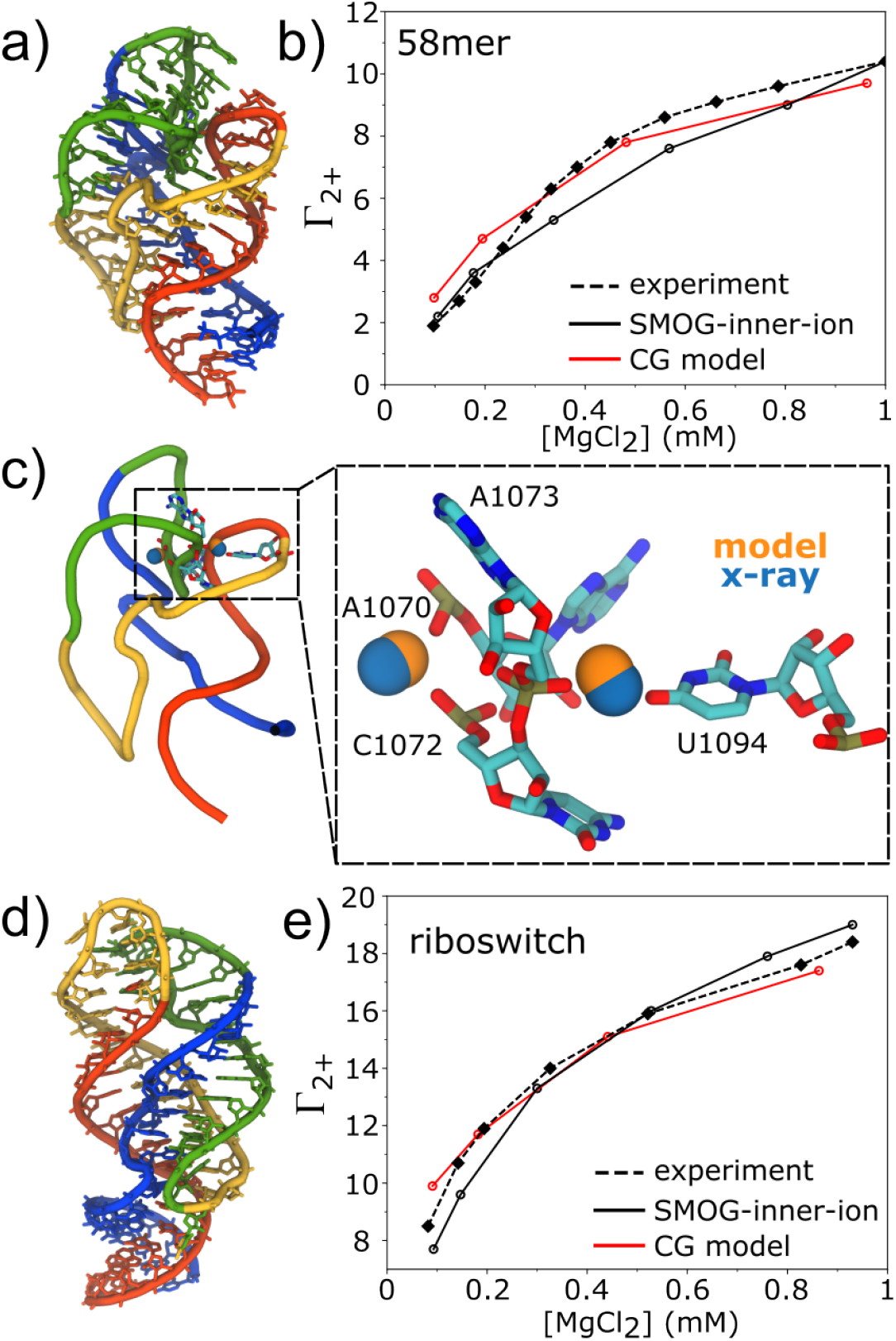
SMOG+inner-ion model accurately describes the concentration-dependent ionic environment and bidentate binding sites. a) Structure of a well-studied 58-mer rRNA fragment (RCSB ID: 1HC8^88^). b) The predicted and experimentally measured preferential interaction coefficient Γ^2+^ for the 58-mer over a range of Mg^2+^ concentrations with [KCl] = 150 mM. The level of agreement is comparable to a leading coarse-grained model with implicit monovalent ions.^50^ c) The model also accurately predicts crystallographically-reported bidentate binding positions. Crystallographic positions are shown as blue spheres, and the average positions in the simulations are shown in orange. Both sites are predicted to within 0.6 Å of the experimental positions. d) Crystallographic structure of the Ade riboswitch. e) Comparison of predicted and experimental Γ^2+^ values for the adenine riboswitch ([KCl] = 50 mM). There is good agreement with the experimental values and those obtained with a coarse-grained model.^50^ Since the Ade riboswitch was not used to parameterize the model, this demonstrates the utility of the model for different RNA systems and monovalent/divalent ionic concentrations.

We find that the SMOG+inner-ion model can accurately predict the dependence of Γ_2+_ on monovalent and divalent ion concentrations. For the 58-mer, the SMOG+inner-ion model yields Γ_2+_ values that agree with previous experimental measurements^86^ (Fig. 2b). At the lowest and highest Mg^2+^ concentrations, Γ_2+_ is within 0.2 of the experimentally determined values, and it is underestimated by approximately 1 at intermediate concentrations. Similarly, the model predicts Γ_2+_ values for the Adenine riboswitch that are within 1 of the experimental values for the considered range of [MgCl_2_]^90^ (Fig. 2e). Interestingly, the model predicts a higher number of inner-shell Mg^2+^ ions than outer-shell (Fig. S4), though both modes exhibit significant concentration dependencies. These simulations were performed with the same [KCl] values used experimentally (50 and 150 mM). Accordingly, the agreement between predicted and experimental Γ_2+_ values for both systems/concentrations shows that the model provides an experimentally-consistent description of the thermodynamic competition between monovalent and divalent ions.

The model also accurately predicts bidentate binding positions that have been implicated by crystallographic analysis of the 58-mer.^88,91^ With this model, Mg^2+^ spontaneously forms bidentate interactions at two specific sites: 1) between the phosphate oxygen of A1073 and the nucleobase O4 atom of U1094 and 2) between the phosphate oxygen atoms of A1070 and C1072 (Fig. 2c). The predicted binding positions of these ions are within 0.6 Å of the crystallographically-resolved sites, highlighting the spatial precision of the model. Further, the A1073/U1094 site is occupied with a probability that exceeds 0.99 ([MgCl_2_] = 1 mM). This is consistent with the high affinity predicted by non-linear Poisson-Boltzmann calculations.^88,92^ For the A1070/C1072 position, the probability of occupancy is also high (~ 0.9), though individual ions interconvert between bidentate and unidentate forms. In the unidentate state, the ion simultaneously forms one inner-shell and one outer-shell interaction (Fig. S5).

### Ribosomal subunit rotation responds to changes in Mg^2+^ concentration

Since the SMOG+inner-ion model can accurately describe ionic interactions in small (PO^−^_4_) and intermediate-scale (58-mer and Ade RNA) systems, we applied it to investigate the role of Mg^2+^ ions during a large-scale conformational rearrangement in the yeast ribosome (Fig. 3a). Specifically, we asked whether intersubunit Mg^2+^-mediated interactions can significantly influence the energetics and kinetics of small subunit (SSU; ~ 1 MDa) rotation, a rearrangement that is required for mRNA-tRNA translocation (i.e. movement of tRNA molecules between ribosomal binding sites during elongation).^93–98^ Rotation is associated with structural rearrangements at the subunit interface, which is composed primarily of RNA-RNA backbone interactions.^99,100^ This raises the possibility that cation-mediated interactions may dynamically reorganize in order to regulate the energetics and kinetics of rotation.

**Figure 3:**
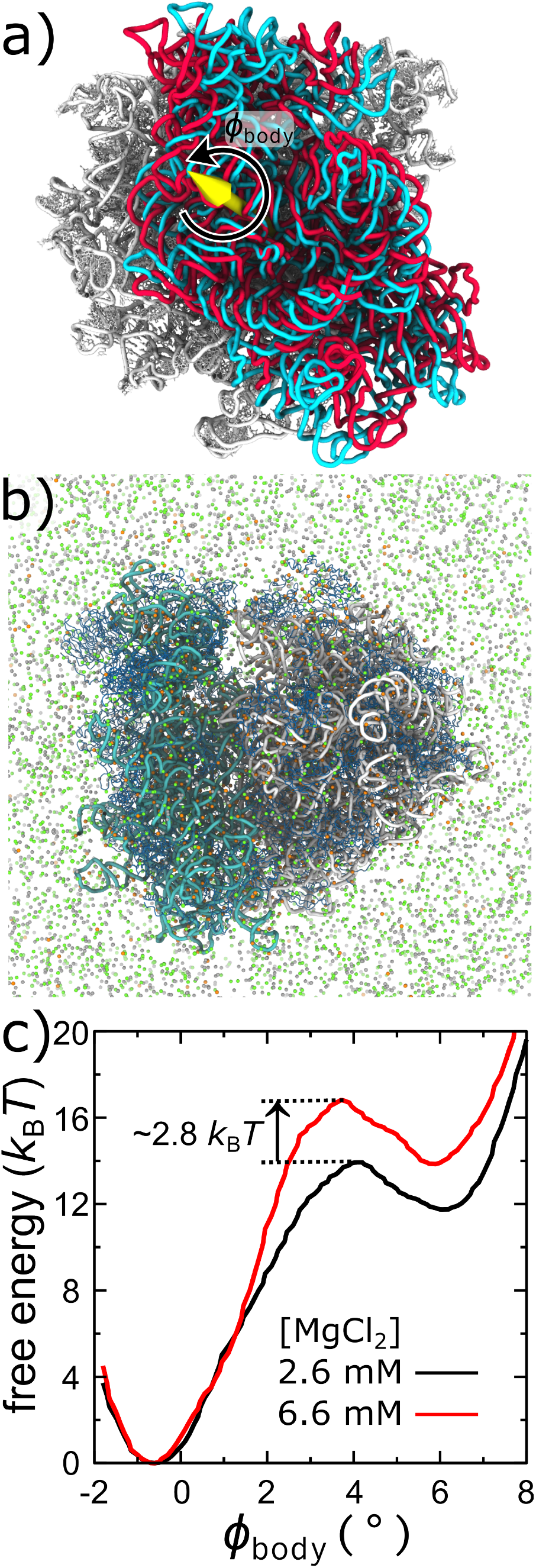
Quantifying the direct impact of Mg^2+^ ions on large-scale dynamics in the yeast ribosome. a) During translation, the small subunit (SSU) of the ribosome must rotate/back-rotate relative to the large subunit (LSU). The 25S rRNA of the LSU is shown in white, with the 18S rRNA of the SSU shown in the unrotated (cyan) and rotated (red) conformations.^77^ The angle *φ*_body_ is used to measure intersubunit rotation.^105^ Representative simulated snapshot of the ribosome (23S, white; 18S, cyan; proteins, blue) with physiological concentrations of Mg^2+^ (orange), K^+^ (green) Cl^−^ (silver) ions. The free-energy of the ribosome as a function of subunit rotation, calculated at [MgCl_2_]=2.6 mM and 6.6 mM. At the higher concentration, there is a shift towards the unrotated ensemble, which is associated with an increase of ~ 2.6*k*_B_*T* in the barrier. This corresponds to a roughly 10-fold increase in the population of the unrotated state and a ~16-fold decrease in the rate of forward rotation.

To identify the roles of ions during subunit rotation, we applied the SMOG+inner-ion model to simulate the complete 80S yeast ribosome.^77^ These simulations included every non-Hydrogen atom in the ribosome, as well as explicit Mg^2+^, Cl^−^ and K^+^ ions (Fig. 3b). In contrast to our simulations of small RNA systems, simulations of the ribosome employed a multi-basin variant^76,101–103^ of the structure-based force field, in which the rotated and unrotated structures are defined to be potential energy minima (See methods). Rotation/back-rotation transitions are two-state when using an electrostatics-free version of the model,^76^ consistent with recent single-molecule measurements of the yeast ribosome.^104^ Here, the SMOG+inner-ion model introduces electrostatic/ionic effects that can shift the energetic balance between endpoints and impact the free-energy barrier. Accordingly, the current objective is to determine if there are site-specific ion-mediated interactions that depend on concentration and contribute to the thermodynamics/kinetics of a large-scale conformational rearrangement.

Since the composition of bound ions on the eukaryotic ribosome has not been unambiguously determined, we performed multiple calculations to establish an initial spatial distribution. First, we performed extensive equilibration simulations, which allowed large numbers of inner-shell Mg^2+^ ions to associate, dissociate and reorganize on the ribosome (Fig. S6a,b). After equilibration, there were approximately 1050 inner-shell and 400 outer-shell ions on the ribosome (Fig. S6c). To provide support for our initial distribution, we tested our model using a 1.55 Å resolution structure of a bacterial ribosome for which strict stereochemical considerations were applied when assigning the ionic composition.^33^ For this experimental structure, we performed energy minimization and found that more than 90% of the ions were displaced by less than 2Å from their experimental positions, which verifies that the energetics in our model are generally consistent with experimental ion assignments for the ribosome.

Simulations with the SMOG+inner-ion model predict that subunit rotation depends significantly on [MgCl_2_], where the unrotated (classical) state is stabilized at higher concentrations. This is reflected in the free energy as a function of SSU rotation (measured by *φ*_body_, defined previously^105^), which shifts towards the unrotated ensemble at [MgCl_2_]=6.6 mM, relative to 2.6 mM (Fig. 3c). Specifically, the free-energy of the unrotated ensemble (*φ*_body_ ~ −0.5^*?*^) decreases by approximately 2 *k*_B_*T* relative to the rotated ensemble (*φ*_body_ ~ 6^*?*^). This predicted change is consistent with single-molecule FRET measurements of a bacterial ribosome, which found the classical state is adopted with higher frequency at higher [MgCl_2_].^106,107^ In contrast to the smFRET studies, which included tRNA molecules, our simulations describe a tRNA-free ribosome. While ion-mediated tRNA-ribosome interactions may also contribute to the energetics of translocation, the current analysis shows that the energetics of rotation directly respond to changes in the ionic conditions.

The predicted concentration-dependent energetics of rotation provides a physical explanation for previous experimental measurements of tRNA translocation kinetics in bacteria. Here, the free-energy barrier associated with forward rotation increases by roughly 2.8 *k*_B_*T* at [MgCl_2_]=6.6mM (Fig. 3c), implicating a ~16-fold reduction in the rate of forward rotation. This is consistent with the observation that the rate of tRNA translocation is ten times slower at [MgCl_2_] of 6mM than 1mM.^108^ Since subunit rotation is required for tRNA translocation to occur,^93^ this suggests that the observed slow down is due to a change in rotation dynamics, which then limits the kinetics of tRNA translocation.

### Dynamics of site-specific ion-mediated interactions during rotation

The simulations reveal examples for how site-specific ion-mediated interactions can contribute to the concentration-dependent energetics of the ribosome. Intersubunit ions are associated with two types of interactions. First, an “inner–outer” ion forms an inner-shell/chelated interaction with one subunit and an outer-shell interaction with the other. Second, an “outer-outer” ion simultaneously forms outer-shell interactions with both sub-units. Inner-inner ions were not found to bridge the subunits. This is consistent with expectations, since bindentate binding interactions could lock the ribosome in a single rotation state, which would stall translation.

Analysis of the simulated dynamics show how concentration-dependent intersubunit interactions introduce energetic competition between rotation states, leading to increased stability of the unrotated ensemble at higher ionic concentrations (Fig. 3c). To identify ion-mediated effects, we first calculated 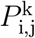: the probability that residue *i* of the LSU and *j* of the SSU simultaneously interact the same Mg^2+^ ion in rotation state *k* (Fig. S7). For this, an Mg^2+^ ion had to be within 5 Å of the backbone oxygen atoms. 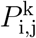 was calculated for [MgCl_2_]=6.6 and 2.6 mM, where the difference 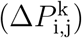 was used to identify subunit-bridging ions that contribute to concentration-dependent rotation energetics. For the unrotated ensemble, the 10 interactions that exhibit the largest increase correspond to four distinct sites (numbered 1-4; Figs. 4a, 5a). A site is composed of all residues that can interact with a single ion. For example, an Mg^2+^ was observed to simultaneously interact with A2259 (LSU) and G1645, C1646 and U1647 (SSU). Accordingly, site 1 is defined by the interactions A2259-G1645, A2259-C1646 and A2259-U1647. Multiple association sites in the rotated state are also responsive to changes in [MgCl_2_] (Fig. 5B). Of these highly-responsive sites, some are partially occupied at both concentrations, whereas others are fully occupied at 6.6 mM, and some are unoccupied at 2.6 mM (Fig. S7). Site 1 is implicated in both the unrotated and rotated states, suggesting that it simply stabilizes the LSU-SSU interface by bridging H69 and h44. In contrast, sites 2, 3 and 4 specifically stabilize the unrotated state at higher concentrations. Sites 5, 6 and 7 have an opposing trend, where they can impart stability to the rotated ensemble at higher ionic concentration. Together, these examples illustrate how ions can introduce competing state-specific contributions to rotation, where the net effect is to stabilize the unrotated state at higher Mg^2+^ concentrations.

**Figure 4:**
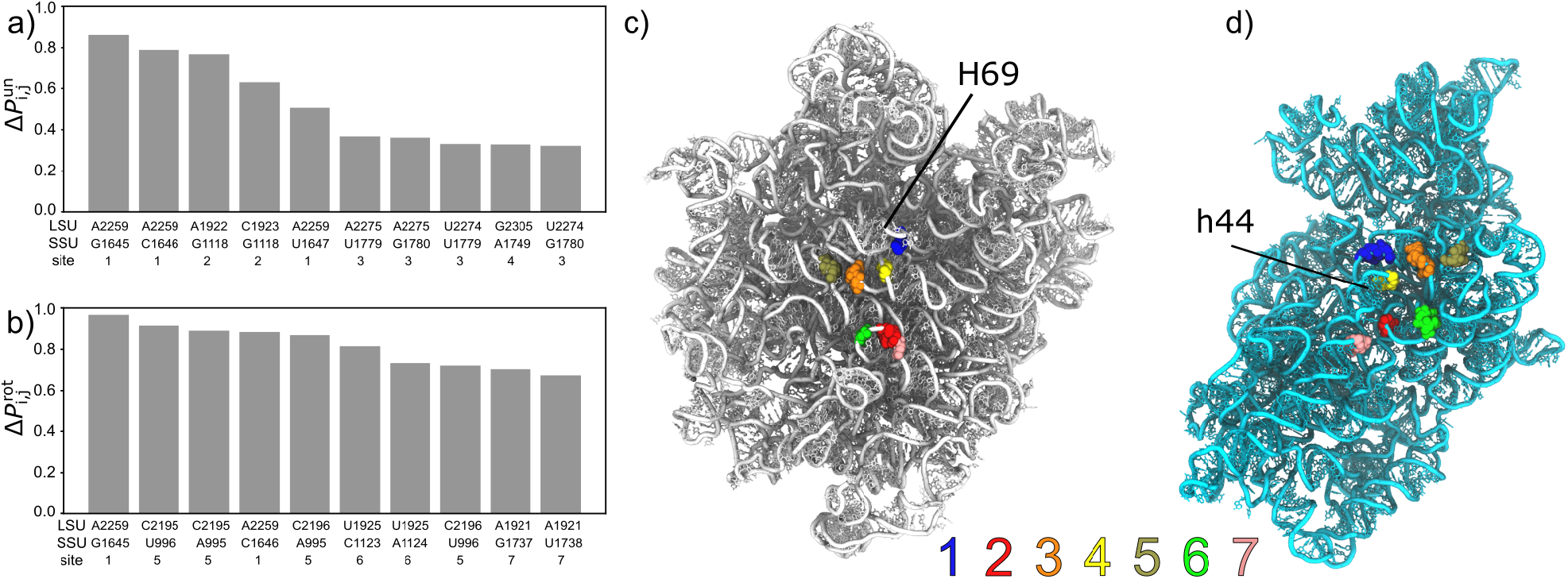
Concentration-dependent intersubunit ion association sites. To identify ion-mediated interactions that contribute to the concentration-dependent energetics of subunit rotation, we calculated 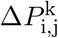: the difference in the probability to form an ion-bridging intersubunit interaction (between residues i and j) at [MgCl_2_]=2.6mM and 6.6 mM for state k (unrotated, panel a; rotated panel b). Calculations were performed with [KCl] = 100 mM. The ten highest 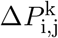 values are shown for both rotation states, where the implicated interactions correspond to seven distinct sites on the intersubunit interface. Of these, site 1 is common to both rotation states, suggesting it contributes to overall affinity of the subunits. In contrast, the response of sites 2, 3 and 4 are specific to the unrotated state and sites 5, 6 and 7 are specific to the rotated state. Structure of the large subunit (panel c) and small subunit (panel d) rRNA, shown from the perspective of the intersubunit interface. Concentration-dependent intersubunit ion sites are labeled.

**Figure 5:**
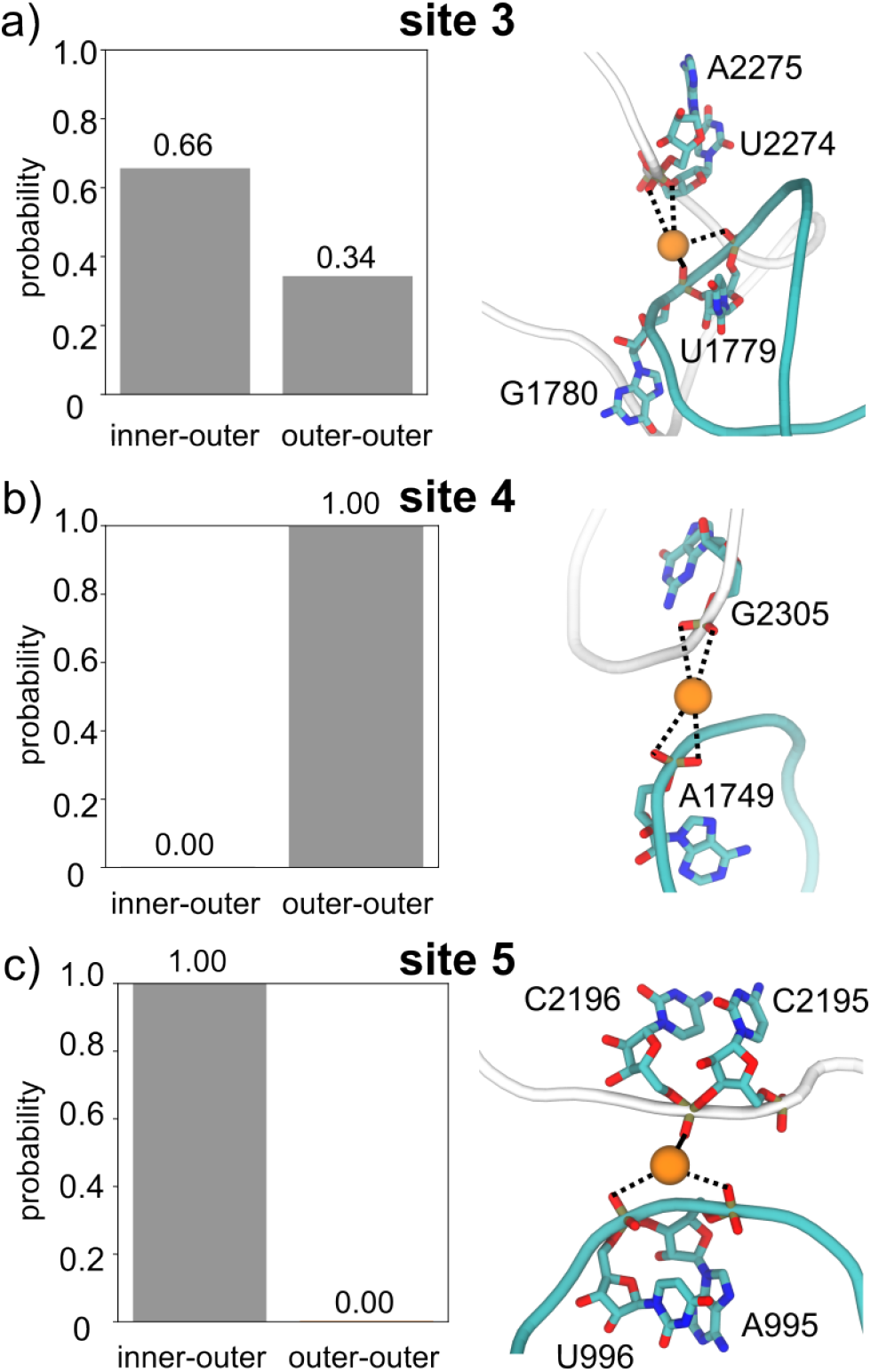
A combination of inner-shell and outer-shell ionic interactions contribute to the energetics of rotation in the ribosome. a) Mg^2+^ ions at site 3 (Fig. 4) are more likely to form an inner-shell interaction with one subunit and an outer-shell interaction with the other (inner-outer), though they often form outer-shell interactions with both subunits (outer-outer). b) Ion-mediated interactions at site 4 were only associated with outer-shell interactions, whereas ions at site 5 (panel c) always formed at least one inner-shell interaction. In the representative structures, dashed lines indicate outer-shell interactions, while inner-shell interactions are shown with solid lines. Statistics for other sites are shown in Fig. S8.

The model shows how a combination of outer-outer and inner-outer interactions can interact with highly dynamic regions of the ribosome in order to regulate the energetics of subunit rotation. We find that ions at sites 1 (Fig. S8a), 5 (Fig. 5c) and 6 (Fig. S8c) only associate in an inner-outer state. In contrast, Mg^2+^ ions at sites 2 (Fig. S8b) and 4 (Fig. 5b) exclusively form outer-outer interactions between subunits. However, sites 3 (Fig. 5a) and 7 (Fig. S8d) are more interesting, where ions can adopt both inner-outer and outer-outer states. This further highlights the dynamic character of the interface, where there is a high degree of molecular flexibility (Fig. S9) that allows ions to interconvert between binding modes. That is, the association sites are not static entities, but are rather constantly undergoing structural fluctuations, where ions can transiently capture these dynamic elements and stabilize the endpoints of rotation.

The dynamic properties of the interface make it difficult to experimentally identify intersubunit-bridging ions. An early crystallographic structure of a bacterial LSU did not detected ions along the peripheral RNA segments, which may be due to lower quality data for these regions and less localized ionic positions.^14^ Another early crystallographic structure of the full ribosome did identify a small number of intersubunit-bridging ions,^18^ and later structural analysis showed that the formation of intersubunit-bridging ions can depend on the conformational state of the ribosome.^19^ In contrast to crystallographic efforts, intersubunit bridging ions were not found when a stereochemical-based approach was applied to identify ion binding sites in a previously-reported 1.55Å cryo-EM structure of a bacterial ribosome.^21,33^ Since association sites at the interface are likely to be populated in a probabilistic and concentration-dependent manner (Fig. S7), the lack of ions in cryo-EM reconstructions is not surprising. Accordingly, one can expect the ions to be less localized, which will lead to weaker experimental densities. Consistent with this argument, the stereochemical analysis only identified ions deep within the structure of the ribosome, where the rRNA is least flexible (Fig. S9). Thus, while modern structural techniques can readily identify highly-localized ions, new methods are required to fully characterize how ions control dynamic processes.

## Discussion

Despite impressive advances in experimental, theoretical and computational capabilities, identifying the precise roles of ions during dynamic processes in RNA has remained elusive. Structural techniques have captured long-lived chelated ions, while other methods have identified the composition of the local ionic environment.^6,109^ In addition, there have been numerous efforts to use highly-detailed explicit-solvent models to characterize short-time dynamics of ions.^39–41,110^ However, biologically-relevant conformational rearrangements occur over time and length scales that are often inaccessible with conventional models. Further, the structural elements involved in functional processes often exhibit a significant degree of disorder, posing major challenges for both experimental and computational methods. In the present study, we showed how one may overcome previous limitations and identify site-specific ionic interactions that provide concentration-dependent contributions to the energetics of large-scale conformational rearrangements.

A range of previous models have been developed to identify the influence of ions on RNA dynamics. In most cases, the models have had utility in the study of folding^50^ and conformational flexibility^25^ in small RNA molecules. A notable example is the coarse-grained model of Nguyen et al,^50^ which can accurately describe folding thermodynamics and has demonstrated cooperativity in the folding of ribosomal RNA.^111^ However, the precise steric content of the ribosome can tightly regulate conformational rearrangements.^112–114^ Accordingly, coarse-grained representations will introduce significant kinetic artifacts for these systems.^53^ Inspired by coarse-grained models with explicit ions,^50,51^ we recently developed an all-atom model that used a simplified energetic scheme for the ribosome^26^ along with explicit Mg^2+^ K^+^ and Cl^−^. While that model describes diffuse ions, it does not account for longer-lived (*µ*s-ms) inner-shell interactions. In the current study, we built upon these earlier efforts and now treat inner-shell interactions of Mg^2+^ and K^+^. By demonstrating the utility of this model to study a conformational rearrangement in the ribosome, the study opens many new questions about the roles and mechanisms by which ions impact large-scale dynamics. We expect that the continued refinement and applications of these models will allow for any number of complex and dynamic assemblies to be studied.

The predicted ionic composition of the ribosome also suggests new strategies for combining experimental and theoretical methods to quantify the ionic environment in large and dynamic assemblies. Experimental methods can readily identify long-lived “structural” ions, though they are not likely to reveal more transient/dynamics interactions. In addition, the stability of inner-shell binding sites can vary significantly, necessitating a probabilistic treatment of ion association events. Since partially-occupied sites will also yield weaker signals, structural data will likely remain ambiguous in these cases. For these dynamic/probabilistic interactions, energetics-based models may now be used to complement and guide interpretation of the data. That is, models can provide strong evidence of inner-shell interactions in dynamic regions, which may then be used to help assign ambiguous cryo-EM densities, or be used to guide crystallographic refinement. Together, the integration of energetic, stereochemical and structural approaches can provide a definitive physical description that bridges short-scale transient ionic interactions with large-scale collective dynamics in biological assemblies.

## Acknowledgement

We thank Dr. Yang Wang and Michael Goldstein for discussions. This work was supported by the National Institutes of Health (grant R35GM153502-01) and the National Science Foundation (grant MCB-1915843). Work at the Center for Theoretical Biological Physics was supported by the National Science Foundation (grant PHY-2019745). We thank AMD for the donation of critical hardware and support resources from its HPC Fund that made this work possible. We also acknowledge generous support from the Northeastern University Explorer cluster and the Northeastern University Research Computing staff.

## Footnotes

† Ion-guided eukaryotic ribosome dynamics

