## Supplementary material for "Transient ion-mediated interactions regulate subunit rotation in a eukaryotic ribosome": SI

### Supplementary Information for: Transient ion-mediated interactions regulate subunit rotation in a eukaryotic ribosome

#### Contents

**1 SI Methods** **S2**
1.1 Parametrization of ionic interactions . . . . . S2
1.1.1 Initial parameter values . . . . . S2
1.1.2 Parameter refinement protocol . . . . . S2
1.1.3 $Mg^{2+}$ parametrization . . . . . S3
1.2 Free energy calculations . . . . . S3
1.3 Preferential interaction coefficient calculations . . . . . S4
1.4 Binding sites probabilities . . . . . S4
1.5 Testing SMOG+inner-ion model for a high-resolution ribosome structure . . . . . S5
1.6 Simulation details . . . . . S5
1.6.1 Explicit-solvent simulations . . . . . S5
1.6.2 SMOG+inner-ion simulations . . . . . S6

**2 SI Results** **S6**
2.1 Disfavoring uni-bidentate binding . . . . . S6
2.2 Robustness of Mg-O dissociation rates . . . . . S7
2.3 Refined effective potential parameters . . . . . S7

#### List of Figures

S1 Disfavoring uni-bidentate interactions . . . . . S8
S2 Diffusion coefficients determine reduced time unit . . . . . S9
S3 Mg-Phosphate lifetime . . . . . S10
S4 Average number of Inner- and Outer-shell $Mg^{2+}$ ions around smaller RNA systems . . . . . S11
S5 $Mg^{2+}$ ions can interconvert between bidentate and unidentate forms . . . . . S12
S6 Equilibration of inner-shell ions on the ribosome . . . . . S13
S7 Concentration-dependent intersubunit ion association sites . . . . . S14
S8 Probability profiles for inner-outer and outer-outer interactions at ribosomal sites . . . . . S15
S9 Structural flexibility of the subunit interface . . . . . S16

### 1 SI Methods

#### 1.1 Parametrization of ionic interactions

The parameters of the effective potential $V_{\mathrm{E}}$ (Eq. 4) were initially parameterized based on explicit-solvent simulations. As reported previously[1], the potential of mean force (PMF) for pairwise interactions was calculated using explicit-solvent molecular dynamics simulations. Radial distribution functions (RDFs) were obtained for various ion–RNA, ion–ion, and ion–protein pairs, and the corresponding PMFs were extracted. These PMFs served as target reference data for fitting the initial form of the effective potential, where an iterative refinement protocol was applied to match the target PMF. In the current study, we build upon these earlier steps in order to parametrize long-lived inner-shell Mg-O interactions. The steps described in sections 1.1.1 and 1.1.2 were performed previously [1], while the current study involved the introduction of steps described in section 1.1.3. For completeness, a full methodological description is provided here.

##### 1.1.1 Initial parameter values

Initial estimates for the parameters were obtained by fitting the potential for pairwise interactions to the PMFs calculated from explicit-solvent simulation of h44 rRNA and ribosomal protein S6 in [$MgCl_2$]= 10 mM and [KCl]= 100 mM. Specifically, the initial values of the effective potential $V_{\mathrm{E}}$ were set by fitting the functional form of the potential (Eq. S1) to the PMF for each type of interaction (*e.g.* Mg-Cl, Mg-O, K-Cl). The Coulomb term was subtracted from the PMF in order to fit the parameters of the ionic excluded-volume term and solvation-shells term $(A, B_{ij}^{(k)}, C_{ij}^{(k)}, R_{ij}^{(k)})$. The values of $A$ in the excluded volume term $V_{\mathrm{ion-excl}}$ were initially set to fit the potential at the shortest distance in the PMF. Additionally, the coefficients of the Gaussians $(B_{ij}^{(k)}, C_{ij}^{(k)}, R_{ij}^{(k)})$ , which account for the solvation shells term $V_{\mathrm{sol}}$, were set to fit the height, depth, width and centers of the peaks and wells in the PMF.

##### 1.1.2 Parameter refinement protocol

The parameters $A$ and $B^{(k)}$ were next refined using the iterative protocol of Savelyev and Papoian[2, 3]. The potential may be expressed as :

$$\begin{aligned} V_{\mathrm{E}} &= V_{\mathrm{coulomb}} + V_{\text{ion-excl}} + V_{\mathrm{sol}} \\ &= \sum_{ij} \frac{q_i q_j}{4\pi\epsilon\epsilon_0 r_{ij}} + \sum_{ij} \frac{A}{r_{ij}^{12}} + \sum_{ij} \left( \sum_{k=1}^{7} B_{ij}^{(k)} e^{-C_{ij}^{(k)}[r_{ij}-R_{ij}^{(k)}]^2} \right) \end{aligned} \tag{S1}$$

With the exception of the Coulomb term, once static values are assigned to $C^{(k)}$ and $R^{(k)}$, the potential can be expressed as:

$$H_{\mathrm{R}} = \sum_{\alpha=1}^{N} K_\alpha S_\alpha, \tag{S2}$$

where the Hamiltonian $H_{\mathrm{R}}$ is expressed as a set of $N$ observables $\{S_\alpha\}$ with weights $\{K_\alpha\}$. These weights correspond to the $\{A, B^{(k)}\}$ coefficients in the potential while $S_\alpha$ corresponds to $\sum_{i<j} \frac{1}{r_{ij}^{12}}$ and $\sum_{i<j} e^{-C^{(k)}[r_{ij}-R^{(k)}]^2}$ for all interaction pairs. As a note, during iterative refinement the $k = 1$ and $k = 2$ (i.e inner-shell terms specific to Mg-O interactions) terms were excluded, such that outer-shell parameters could be obtained.

With this linear functional form of the potential, the parameters could be iteratively refined, using the observable values calculated from explicit-solvent simulations as a target. Let the expectation value of the observable $S_\alpha$ be given by $\langle S_\alpha \rangle_{\mathrm{ion-model}}$, calculated from a simulation performed using a structure-based model with ions and $\langle S_\alpha \rangle_{\mathrm{ex-sol}}$, calculated from explicit-solvent simulations. To identify the values of $K_\alpha$ that best reproduce the target observables, one may Taylor expand $\langle S_\alpha \rangle_{\mathrm{ion-model}}$ with respect to $K_\alpha$, about an initial set of weights $\{K_\alpha^{(0)}\}$. To first order, the difference between the simulated and reference expectation values ($\Delta\langle S_\alpha\rangle = \langle S_\alpha\rangle_{\mathrm{ion-model}} - \langle S_\alpha\rangle_{\mathrm{ex-sol}}$) is given by: [2, 3]:

$$\Delta\langle S_\alpha\rangle = \sum_\gamma \frac{\partial\langle S_\alpha\rangle}{\partial K_\gamma}\Delta K_\gamma \tag{S3a}$$

$$= \frac{1}{k_B T}\sum_\gamma [\langle S_\alpha S_\gamma\rangle - \langle S_\alpha\rangle\langle S_\gamma\rangle]\Delta K_\gamma, \tag{S3b}$$

where $\Delta K_\gamma^{(n)} = K_\gamma^{(n+1)} - K_\gamma^{(n)}$.

To obtain ion parameters, simulations with the SMOG+inner-ion model were performed in each iteration, followed by an update of the weights, where in the $n$th iteration, the new weight is defined as $K_\alpha^{(n+1)} = K_\alpha^{(n)} + \Delta K_\alpha^{(n)}$. Accordingly, the final values of the coefficients $A$ and $B_{ij}^{(k)}$ for the SMOG+inner-ion model are extracted from the converged $K_\alpha$ values. This provided a set of parameters that describe the outer-shell interactions in the model.

##### 1.1.3 $Mg^{2+}$ parametrization

To account for inner-shell interactions, we then modified the $A$ values (excluded volume), as well as the $k = 1$ and $k = 2$ (inner-shell and barrier) parameters. The specific values were manually adjusted for Mg-O interactions, in order to account for the proper coordination distance, binding affinity and dissociation rate. As described in the main text, the target values were based on a combination of previous experimental observations and ab-initio calculations. Finally, the weight of the outer-shell interaction ($B_{ij}^{(3)}$) was adjusted based on estimates obtained from free-energy perturbation calculations that were used to match the preferential interaction coefficient for the 58-mer rRNA[4] at a single set of ionic conditions: [$MgCl_2$] = 1 mM, [KCl]=150 mM.

#### 1.2 Free energy calculations

To produce the free-energy profiles that were used to describe the energetics of reference compounds and the ribosome, we used umbrella sampling[5]. In the parameterization process, a phosphate group was used as a reference compound. For this system, we used the distance $r$ between a phosphate oxygen ($O_P$) and the $Mg^{2+}$ ion as the umbrella coordinate for the one-dimensional profile (Fig. 1d). The umbrella was given by a harmonic potential centered at $r_0$. The details of the distances and force constants are shown in Table S1.

**Table S1:** Umbrella sampling windows and associated force constants

| **Coordinate** | **Window Range** [Å] | **Force Constant** $k$ [ru/nm$^2$] |
|---|---|---|
| $r_0$ (spacing = 0.05 Å) | 1.50 – 1.85 | 250000 |
| | 1.90 – 2.25 | 100000 |
| | 2.30 – 2.65 | 250000 |
| | 2.70 – 3.45 | 80000 |
| | 3.50 – 3.85 | 20000 |
| | 3.90 – 4.25 | 10000 |
| $r_0$ (spacing = 0.1 Å) | 4.30 – 8.00 | 10000 |

For the two-dimensional profile in Fig. S1, we used $r_0$ as the first coordinate and the coordination number ($CN$) of the $Mg^{2+}$ ion as the second coordinate. $CN$ is defined as:

$$CN = \sum_{i=1}^{3}\frac{1}{2}\{1 - \tanh[\gamma(r_i - \xi)]\}, \tag{S4}$$

where $\gamma = 25$, $\xi = 2.7$Å, and $r_i$ is the distance between the $Mg^{2+}$ ion and the other three phosphate oxygen atoms. 64 umbrella windows were used for the first coordinate $r_0 = [1.50 - 5.00$ Å$]$ at each $CN$ value. For the second umbrella coordinate $CN$, we used a spacing of 0.1 between windows and a harmonic force constant $k = 500\epsilon$.

To generate the free-energy landscape for rotation of the ribosomal subunit, $\Delta Q$ was used as an umbrella coordinate. $\Delta Q$ is the difference between the normalized unique native contacts formed in the rotated and unrotated conformations, and it is defined as:

$$\Delta Q = Q_{\text{rot}} - Q_{\text{unrot}} \tag{S5}$$

where

$$Q_s = \frac{1}{N_s} \sum_{j \in \{C_s\}} \frac{1}{1 + e^{\beta(r_j - \mu r^0_{j,s})}}. \tag{S6}$$

$\{C_s\}$ is the set of unique native contacts in conformation $s$ (rotated, or unrotated), $r^0_{j,s}$ is defined as the distance of contact $j$ in conformation $s$, $\beta = 10$ nm$^{-1}$ and $\mu = 1.8$. A force constant of $700\epsilon$ was used for all umbrella windows with $\Delta Q$ spacing of 0.1 from -0.8 to 0.7. The openmm-plumed[6, 7] package was used to introduce the $\Delta Q$ umbrella potential. Then, we calculated the free-energy profile as a function of the reaction coordinate $\phi_{body}$, which describes rotation of the small subunit relative to the large subunit (defined previously [8]; fig. 3a). $\phi_{body}$ has been demonstrated to be effective in characterizing the underlying energy barrier as well as reflecting diffusive dynamics for rotation in the yeast ribosome[9] . The free-energy profiles were calculated using the Weighted Histogram Analysis Method[10].

#### 1.3 Preferential interaction coefficient calculations

$\Gamma_{2+}$ represents the excess $Mg^{2+}$ ions around the RNA at bulk density ($\rho_{\text{Mg}^{2+}}$). $\Gamma_{2+}$ can be calculated as the difference between the total number of $Mg^{2+}$ ions ($N_{\text{Mg}^{2+}}$) and the value that would correspond to the bulk: $\Gamma_{2+} = N_{\text{Mg}^{2+}} - \rho_{\text{Mg}^{2+}} \times V_{\text{box}}$, where the bulk number is the bulk density times the volume of the box. To calculate $\rho_{\text{Mg}^{2+}}$, the simulated box was partitioned into five equal-width cross-sectional cuboids, after centering the system such that the RNA was in the central cuboid. The density was calculated from the average number of $Mg^{2+}$ ions in the four other cuboids. $\Gamma_{2+}$ was calculated over a range of $Mg^{2+}$ concentration of (0.1 - 1 mM) for the 58-mer and adenine riboswitch. Due to the sensitivity of the bulk density to the convergence of the statistics of association and dissociation of ions, we accelerated the sampling of the association and dissociation events of the $Mg^{2+}$ ions by reducing the barrier height ($B^{(2)}_{ij}$ from 6.7 to 4.5) between inner- and outer shells. The barrier width was also adjusted to minimize the effects on the inner and outer shells ($C^{(2)}_{ij}$ from 300 to 200). To calculate $\Gamma_{2+}$, each simulated frame was re-weighted to calculate the value corresponding to the original potential:

$$\begin{aligned}
\Gamma_{2+} &= \langle N_{\text{Mg}^{2+}} - \rho_{\text{Mg}^{2+}} \times V_{\text{box}} \rangle \\
&= N_{\text{Mg}^{2+}} - \langle \rho_{\text{Mg}^{2+}} \rangle \times V_{\text{box}} \\
&= N_{\text{Mg}^{2+}} - \frac{\sum \rho^{\text{lb}}_{\text{Mg}^{2+}} \times \exp(-\beta \Delta U)}{\sum \exp(-\beta \Delta U)} \times V_{\text{box}},
\end{aligned} \tag{S7}$$

where $\Delta U = U_{\text{ob}} - U_{\text{lb}}$. lb indicates the lower barrier potential and ob is the original barrier.

At each specific $Mg^{2+}$ concentration, $\Gamma_{2+}$ was calculated as an average over 8 replicate simulations of duration $2 \times 10^9$ time steps, each. To ensure reliability of the strategy, we compared the $\Gamma_{2+}$ values obtained using this method with values obtained from simulations using the original barrier with [$MgCl_2$] = 1 mM. For 58-mer RNA, the value calculated from the original barrier simulations is $\Gamma_{2+} = 10.1 \pm 1$ compared to 10.4 for the lower barrier re-weighted simulations. Additionally, for adenine riboswitch, it was $19.2 \pm 1$ for original barrier simulations compared to 18.9 for lower-barrier simulations.

#### 1.4 Binding sites probabilities

To calculate the binding-site probabilities, we analyzed the original-barrier simulations and computed the fraction of frames in which each highly negatively charged oxygen atom of the 58-mer engaged in an inner-shell interaction with a $Mg^{2+}$ ion. An interaction is considered inner-shell if the distance between the oxygen atom and the $Mg^{2+}$ ion is < 2.7 Å (i.e. the center of the barrier). Only two sites—A1073–U1094 and A1070–C1072—exhibited probabilities greater than 0.5. To calculate the average position as shown in Fig. 2b, we aligned the RNA in each trajectory frame to the initial crystal structure and then calculated the average position of the $Mg^{2+}$ ion whenever it was in contact with the RNA oxygen atom. We used the chain ID: D of the PDB ID: 1HC8 in the ion position comparison in Fig. 2b.

#### 1.5 Testing SMOG+inner-ion model for a high-resolution ribosome structure

To verify that the SMOG+inner-ion energetics are consistent with ion binding positions that have been identified experimentally for a 1.55 Å resolution structure of the bacterial ribosome[11], we performed energy minimization on the ions in the structure using our model. We used SMOG 2 (v2.6-beta) [12] when generating the force field files. We then performed the energy minimization for the ions while keeping the non-ionic atoms fixed spatially. L-BFGS algorithm in OpenMM [6] was used to perform minimization.

#### 1.6 Simulation details

##### 1.6.1 Explicit-solvent simulations

As described previously, [1], explicit-solvent simulations were used in the initial parameterization of the SMOG+inner-ion model. Simulations were performed using Gromacs v5.1.4[13]. The simulations were performed using the Amber99sb-ildn force field,[14] where the model systems were solvated with SPC/E water molecules[15]. Monovalent ion parameters by Joung and Cheatham [16] and divalent ion parameters by Åqvist[17] were used. The leap-frog integrator was used with Particle-mesh Ewald summation and a cutoff radius of 10 Å. Three systems were simulated to parameterize the ion-ion, ion-RNA and ion-protein interaction parameters.

1. *Ions in solution.* A total of 60 $K^{+}$, 6 $Mg^{2+}$, and 72 $Cl^{-}$, along with 32,635 water molecules, were placed in a 10 nm cubic simulation box to achieve target concentrations of 10 mM [$MgCl_2$] and 100 mM [KCl]. Each system was initially energy-minimized using the steepest descent algorithm, followed by the conjugate gradient method. Equilibration was performed for 1ns at 300K and 1bar using the Berendsen thermostat and barostat[18]. Subsequently, a 1$\mu$s production simulation was conducted using the Nosé–Hoover thermostat[19, 20] and the Parrinello–Rahman barostat [21] .

2. *RNA in ionic solution.* A fragment of 16S ribosomal RNA corresponding to helix 44 (h44), extracted from PDB ID 4V6F[22], was positioned at the center of a rectangular simulation box with dimensions $10 \times 12.2 \times 12.2$ nm$^3$. The long axis of h44 was aligned along the x-axis of the box. To yield bulk concentrations of 10mM [$MgCl_2$] and 100mM [KCl], the system was solvated with 123 $K^{+}$, 14 $Mg^{2+}$, 108 $Cl^{-}$and 47,917 water molecules. To prevent effects of structural deformation of h44 on ion-RNA interactions, all RNA atoms were held fixed at their initial positions using harmonic restraints with a force constant of 1000kJ/mol/nm$^2$. The system was initially relaxed through a two-step energy minimization protocol, beginning with steepest descent minimization, followed by conjugate gradient minimization. This was followed by thermal and pressure equilibration at 300K and 1bar using Berendsen thermostat and barostat. After 20 ns of equilibration, a 1 $\mu$s production simulation was performed with the Nose-Hoover thermostat and Parrinello-Rahman barostat. The initial 20ns of the trajectory, corresponding to the ionic equilibration phase around h44, was excluded from radial distribution function analysis.

3. *Protein in ionic solution.* Protein S6 of a bacterial ribosome (PDB: 4V6F) was isolated and positioned in a rectangular box with dimensions $10 \times 16 \times 16$nm$^3$. To yield a bulk concentrations of 10 mM [$MgCl_2$] and 100 mM of [KCl], 15$Mg^{2+}$, 154 $K^{+}$, 185 $Cl^{-}$and 88,364 water molecules were added to the box. With restraining the positions of the protein atoms to their initial positions with force constant 1000 kJ/mol/nm$^2$, we minimized the energy of the system and run the simulation with the same protocol mentioned earlier with the RNA in ionic solution.

##### 1.6.2 SMOG+inner-ion simulations

All SMOG+inner-ion simulations were performed at temperature of 0.5 reduced units which corresponds to approximately 300 K [23]. All force field were generated using SMOG 2 (v2.6-beta) [12] and simulations were performed using OpenSMOG v1.2 [24]. Upon publication, the SMOG+inner-ion force field files will be available on the smog-server.org force field repository, under accession name AA_inner-ions_Wanes25.v1.

1. *SMOG+inner-ion simulation for a model compound.* During the model refinement process and the calculation of the 1D and 2D free-energy profiles of the Mg–O distance and $CN$, the simulations included a phosphate group, a $Mg^{2+}$ion, and a $Cl^{-}$ion in a cubic box of length 8 nm. For each umbrella window, 234 million time steps were performed. A total of 96 and 704 umbrella windows were used for calculating the 1D and 2D free-energy profiles, respectively.

2. *SMOG+inner-ion simulation for RNA molecules.* In order to caclulate the preferential interaction coefficient $\Gamma_{2+}$ for both 58-mer rRNA (PDB ID: 1hc8) and the adenine riboswitch (PDB ID: 1y26) and calculate the probability of $Mg^{2+}$ binding to sites in the 58-mer, we performed simulations for both systems by placing each at a cube of length 70 nm. For the 58-mer, to yield $Mg^{2+}$ bulk concentration between ($\approx 0.1 - 1$mM), we placed 24 to 217 $Mg^{2+}$ions. Additionally, we added 30,983 $K^{+}$ions to the box for all simulations to achieve $K^{+}$concentration of 150mM. At each $Mg^{2+}$concentration, we performed 8 replicas of simulations with more than $1.7 \times 10^9$ steps each. For the adenine riboswitch, to reach the same $Mg^{2+}$concentration range, 27 to 211 $Mg^{2+}$ions were added. 10,328 $K^{+}$ions were added to set $K^{+}$concentration of 50 mM. For the adenine riboswitch at each $Mg^{2+}$concentration, 8 replicas of the system were simulated for $3.8 \times 10^9$ time step each. For all simulations, $Cl^{-}$were added to neutralize the box charge.

3. *SMOG+inner-ion simulation for Ribosome.* The simulations for 80S ribosome were performed using structures (PDB IDs: 3j77 , 3j78). The rotated and unrotated states are defined to be potential energy minima. To create $K^{+}$concentration of 100 mM, 22,000 $K^{+}$ions were added to the box of length 70 nm. Adding 2000 and 3000 $Mg^{2+}$to the box yields to concentration of 2.6 and 6.6 mM respectively. $Cl^{-}$ions were added to neutralize the box. Since umbrella sampling was used in the ribosome simulations to calculate the PMF of the subunit rotation, 16 umbrella windows were used for each concentration. At each umbrella window, the system was simulated with more than $1.8 \times 10^8$ time steps leading to more than $2.88 \times 10^9$ steps for each $Mg^{2+}$concentration. For the placement of $Mg^{2+}$ ions in the ribosomal system, we employed an in-house tool that first discretizes the volume and assigns an ion to each phosphate oxygen. These preliminary ion positions were then subjected to an energy minimization independently using the SMOG+inner-ion force field with a pairwise cut-off of 2 nm. Following minimization, we retained only those ions exhibiting negative interaction energies and separated by at least 6 Å from one another. This procedure yielded 1,256 uniquely placed $Mg^{2+}$ ions. Any remaining ions required to achieve the desired ionic concentration were added using the smog_ions utility in SMOG2 [12].

### 2 SI Results

#### 2.1 Disfavoring uni-bidentate binding

The uni-bidentate (UBI) conformation (*i.e.* $Mg^{2+}$ ion coordinated with two phosphate oxygen atoms on the same phosphate group) has been shown using ab-initio calculations [25, 26] to have a prohibitively high energy, compared to the unidentate or bidentate states. This is consistent with the lack of observation of this state in high-resolution structures[27]. An explanation[27] based on molecular orbital theory was proposed to interpret the Mg-Phosphate group binding asymmetry suggests that the hybridization of the non-bridging oxygen atoms is a superposition of $sp$ and $sp^2$. In order to make this state unstable in the model, the excluded volume for the Mg-P interaction was adjusted so that the distance between phosphorus and magnesium ion in the chelation state is 3.6 Å, consistent with the value measured in X-ray experiment[28] for Mg-Phosphate complexes. We ensured that the model energetically disfavors this state by calculating the free energy profile of the $Mg^{2+}$ ion approaching two oxygen atoms from the same phosphate group. The free energy profile showed a $> 20$ $k_{\mathrm{B}}T$ difference between this state and the unidentate state(Fig. S1).

#### 2.2 Robustness of Mg-O dissociation rates

To assess the robustness of the calculated dissociation rate, we employed two complementary methods. First, we extracted the rate from the measured half-life (Fig. S3) and converting from reduced time units to effective timescales (See main text). Second, we applied Kramers' theory of barrier crossing[29] to the 19 $k_\mathrm{B}T$ free-energy barrier (Fig. 1d) between the inner- and outer-shell, yielding an estimated rate of $\approx 10^3\mathrm{s}^{-1}$.

#### 2.3 Refined effective potential parameters

As outlined in the Methods section, the SMOG+inner-ion model parameters were initially calibrated against explicit-solvent simulations. Subsequently, a subset of parameters governing ion–RNA interactions was fine-tuned based on comparisons with experimental data. The resulting parameter set is similar to the set described by Wang [1], with additional modifications as detailed in the tables below.

**Table S2:** Refined Parameters for Ion–RNA Interactions. RNA atoms are categorized based on their element type and partial charge. Negatively charged atoms are further grouped according to their element and the magnitude of their partial charge. Atoms with partial charges less than -0.49e are labeled with subscript marked as"<-0.5". In the table, "0" denote that the corresponding interaction parameters are explicitly set to zero, whereas"-" that these parameters are not defined within the SMOG+inner-ion model.

| Interaction | A [$\epsilon \cdot \mathrm{nm}^{12}$] | B [$\epsilon$] | | | | | | |
|---|---|---|---|---|---|---|---|---|
| | A | $B_1$ | $B_2$ | $B_3$ | $B_4$ | $B_5$ | $B_6$ | $B_7$ |
| $K^+$-$O_{<-0.5}$ | $2.923 \times 10^{-8}$ | 0 | 0 | -0.8796 | 0.2962 | 0.0339 | -0.0217 | -0.0973 |
| $Mg^{2+}$-$O_{<-0.5}$ | $8.522 \times 10^{-10}$ | -4.5 | 6.7 | -0.77 | 0.0157 | -0.1161 | 0.0093 | -0.0599 |
| $Mg^{2+}$-P | $4.66 \times 10^{-6}$ | - | - | - | - | - | - | - |

| Interaction | C [$\mathrm{nm}^{-2}$] | | | | | | | R [nm] | | | | | | |
|---|---|---|---|---|---|---|---|---|---|---|---|---|---|---|
| | $C_1$ | $C_2$ | $C_3$ | $C_4$ | $C_5$ | $C_6$ | $C_6$ | $R_1$ | $R_2$ | $R_3$ | $R_4$ | $R_5$ | $R_6$ | $R_7$ |
| $K^+$-$O_{<-0.5}$ | 0 | 0 | 1095 | 501 | 438 | 1833 | 123 | 0 | 0 | 0.278 | 0.348 | 0.511 | 0.569 | 0.698 |
| $Mg^{2+}$-$O_{<-0.5}$ | 800 | 300 | 651 | 392 | 331 | 305 | 92.7 | 0.218 | 0.27 | 0.417 | 0.478 | 0.61 | 0.725 | 1.01 |
| $Mg^{2+}$-P | - | - | - | - | - | - | - | - | - | - | - | - | - | - |

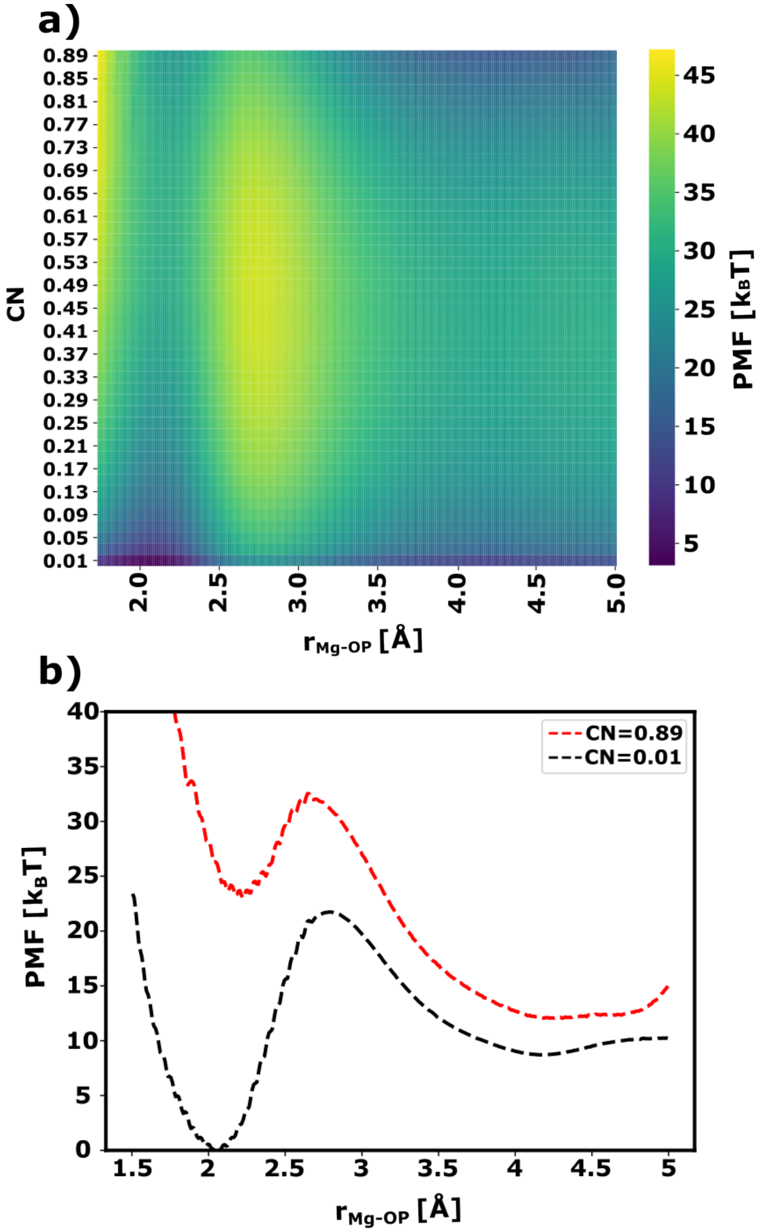


**Figure S1: Disfavoring uni-bidentate interactions** (a) 2D PMF with the distance between $Mg^{2+}$ ion and a phosphate oxygen as the first coordinate and the coordination number $CN$ as the second coordinate. $CN$ is defined to be the number of phosphate oxygen atoms in the inner-shell of the $Mg^{2+}$ ion other than the oxygen atom in the first coordinate (Eq. S4). (b) Projection of $CN$ at values 0.01 and 0.89. This figure shows that when one oxygen atom from a phosphate group forms an inner-shell interaction with a $Mg^{2+}$ ion, it is energetically unfavorable for a second oxygen atom from the same group to simultaneously occupy the inner shell, where the free-energy difference between the unidentate (i.e. one oxygen atom in inner-shell interaction) and uni-bidentate (i.e. two oxygen atoms from the same phosphate group in inner-shell interaction) exceeds 20 $k_{\mathrm{B}}T$.

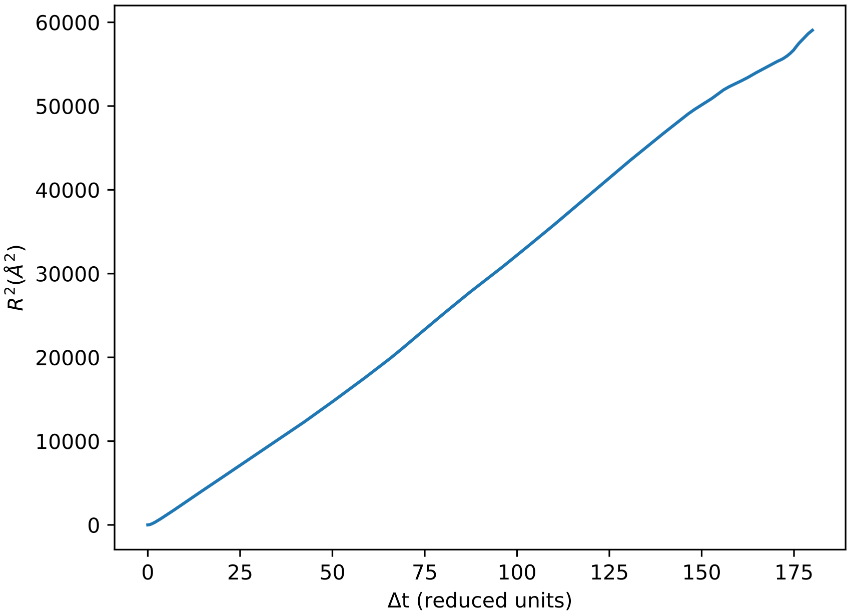


**Figure S2: Simulated time reduced unit determined from diffusion coefficient comparison** The figure shows the displacement-squared of $Mg^{2+}$ ion in a $MgCl_2$ solution as a function of lag time ($\Delta$t) in reduced time unit. Diffusion of $Mg^{2+}$ was found to be approximately 47 $\AA^2/\tau^{eff}$. Comparing the simulation value with the experimental one of $D_0 = 0.71 \times 10^{-5}$ $cm^2/s$[30] indicates that the effective simulated time unit $\tau^{eff}$ is approximately 66 ps.

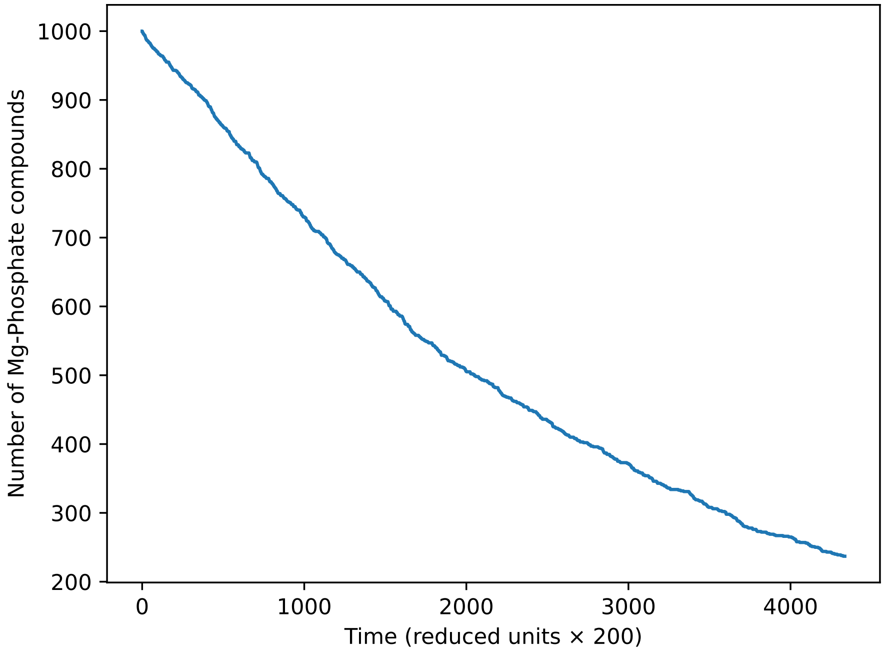


**Figure S3: Determining the lifetime of an inner-shell $Mg^{2+}$ ion with a phosphate oxygen** The mean lifetime was found to be $\approx 0.4 \times 10^{-3}$s by fitting the number of the intact compounds over time as an exponential decay. The mean lifetime value leads to dissociation rate of $2.5 \times 10^{3}s^{-1}$, which lies in the experimentally measured range of $\approx 10^{3} - 10^{4}s^{-1}$[31, 32, 33, 34, 35].

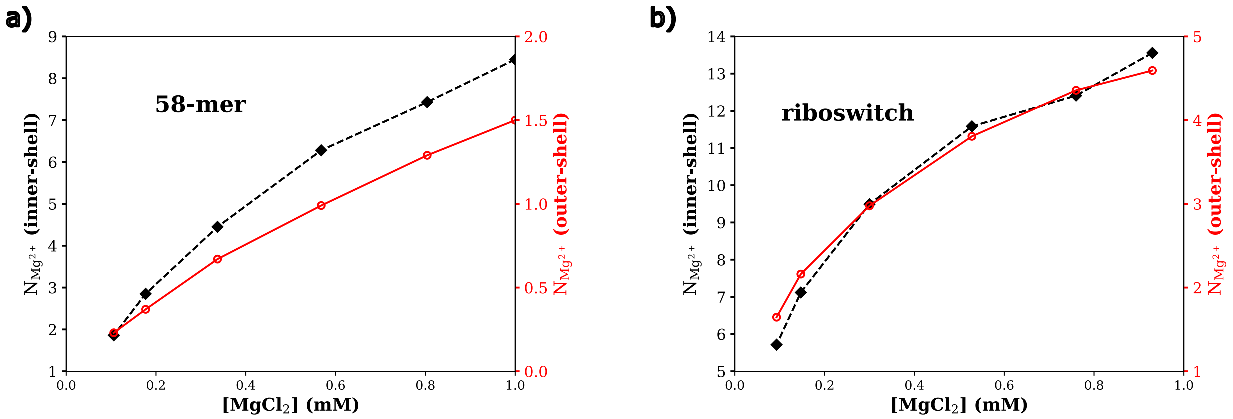


**Figure S4:** a) Average number of inner-shell (black) and outer-shell (red) $Mg^{2+}$ ions associated with the 58-mer at [KCl] = 150mM and over a range of $MgCl_2$ concentration. b) Average number of inner- and outer-shell ions for the adenine riboswitch at [KCl] = 50mM and a range of [$MgCl_2$]. Both RNA systems exhibit monotonic increases in ion association with increasing $Mg^{2+}$ concentration for both inner-shell and outer-shell binding modes, where there is a larger number of inner-shell than outer-shell ions.

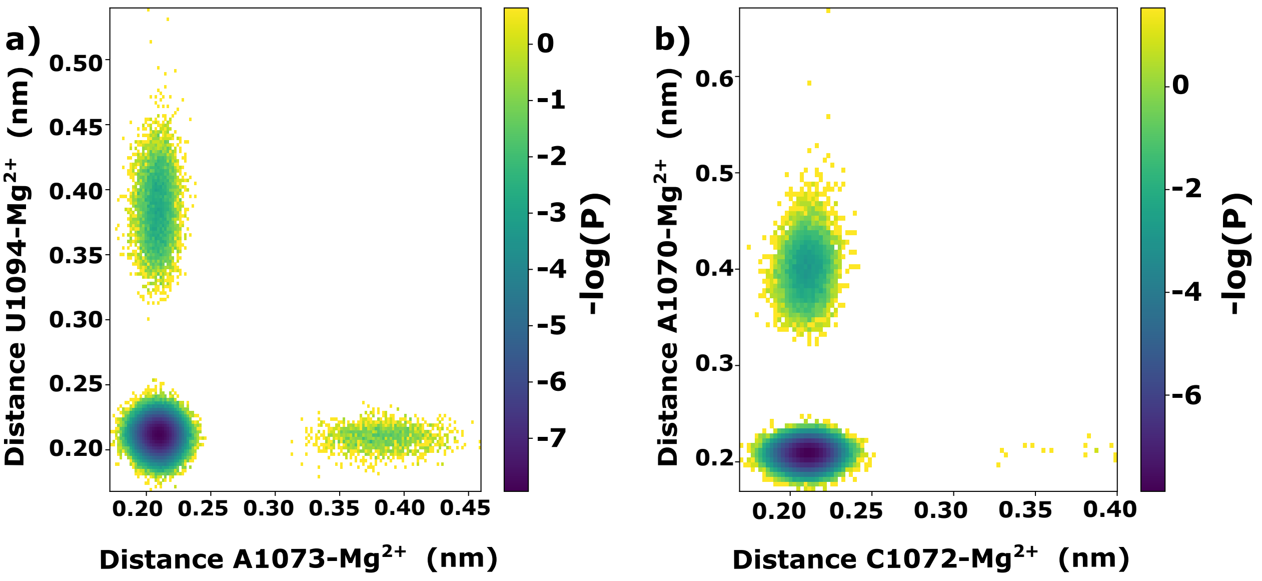


**Figure S5:** $Mg^{2+}$ ions can interconvert between bidentate and unidentate forms within a 58-mer binding site. The figure shows the Negative Log Probability as a function of distances between the $Mg^{2+}$ ion and RNA oxygen atoms of 58-mer at the two most probable positions. The $Mg^{2+}$ ion can form an inner-shell interaction with one oxygen and outer-shell with another oxygen then stabilize by adopting the bidentate state. a) In the first site, the $Mg^{2+}$ ion interacts with A1073-U1094 with comparable probability to be inner-shell with one residue and outer-shell with the other, reflecting a semi-symmetric interaction. b) For the second position where the ion interacts with A1070-C1072, there is a skewed probability profile.

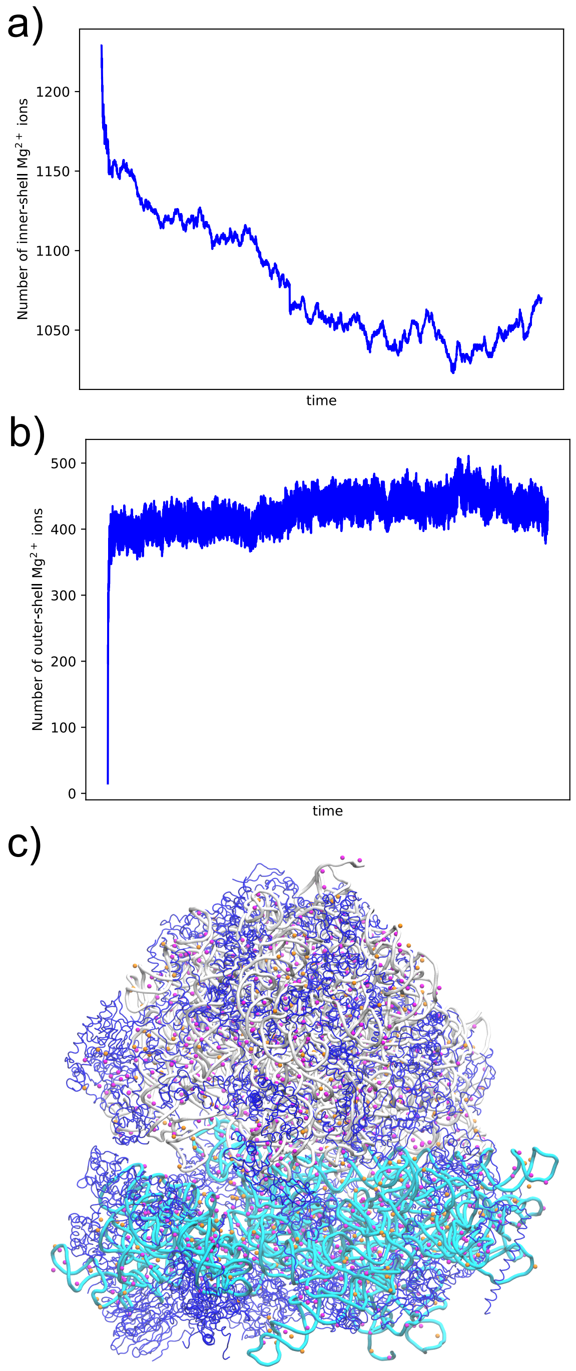


**Figure S6:** After using an energetics-based methods for assigning initial positions for possible inner-shell $Mg^{2+}$ ions, the system was simulated at constant temperature to allow for the ions to equilibrate. During equilibration, the ions freely dissociate, associate or reorganize on the ribosome. a) The number of inner-shell $Mg^{2+}$ ions around non-bridging phosphate oxygen atoms, as a function of simulated time. While the system was initialized with 1256 ions in the inner-shell, the system equilibrated to a state where there were approximately 1050 in the inner shell. b) The number outer-shell $Mg^{2+}$ ions around non-bridging phosphate oxygen atoms, as a function of simulated time. During equilibration, ions transition from the inner-shell positions and bulk, in order to populate the outer-shell around the RNA backbone. c) Representative structural snapshot of the equilibrated system, with inner-shell ions shown in magenta and outer-shell shown in orange.

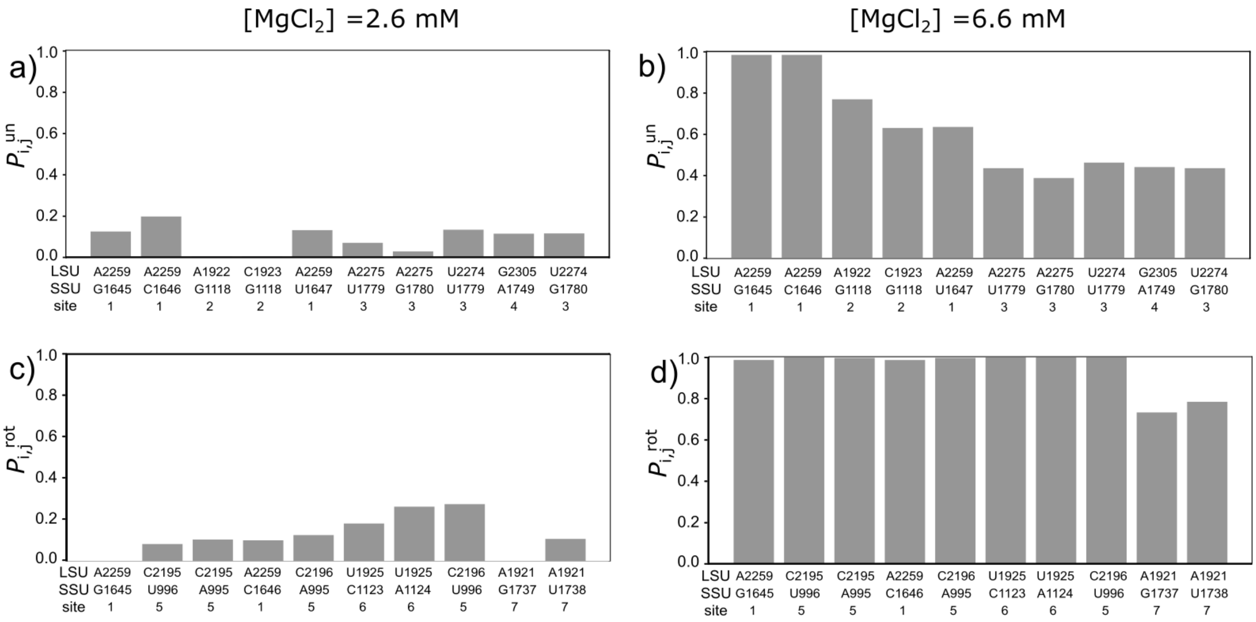


**Figure S7:** To identify the ion-mediated interactions that are the most sensitive to the change to the ionic concentration, we calculated $P^{k}_{i,j}$: the probability to form an ion-bridging intersubunit interaction (between residues i and j) at [$MgCl_2$]=2.6mM and 6.6 mM for state k and calculated the difference between the two concentration as in Fig. 4 a,b. Calculations were performed with [KCl] = 100 mM. Panels a,b represent the probability for the unrotated state at [$MgCl_2$]=2.6 mM and 6.6mM respectively. Panels c,d show the probability for the two concentrations in the rotated ensemble.

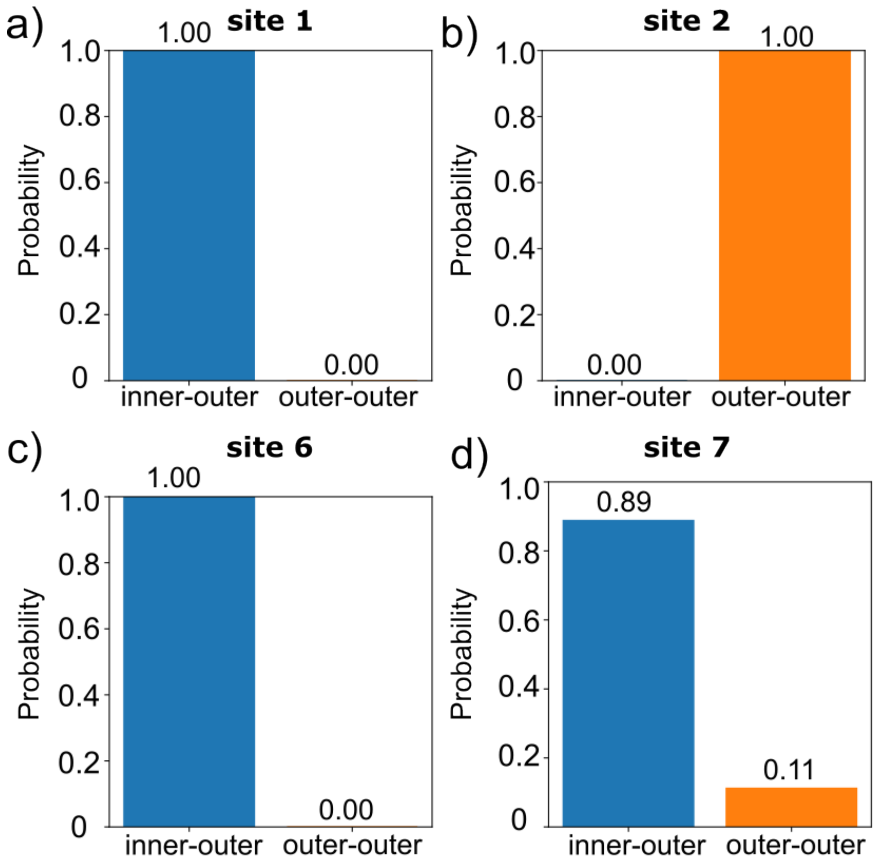


**Figure S8:** Probability profiles for inner-outer and outer-outer interactions at sites 1, 2, 6, and 7 (Fig. 5). The results demonstrate site-dependent heterogeneity in interaction patterns, ranging from purely inner-outer or outer-outer interactions to mixed interaction modes.

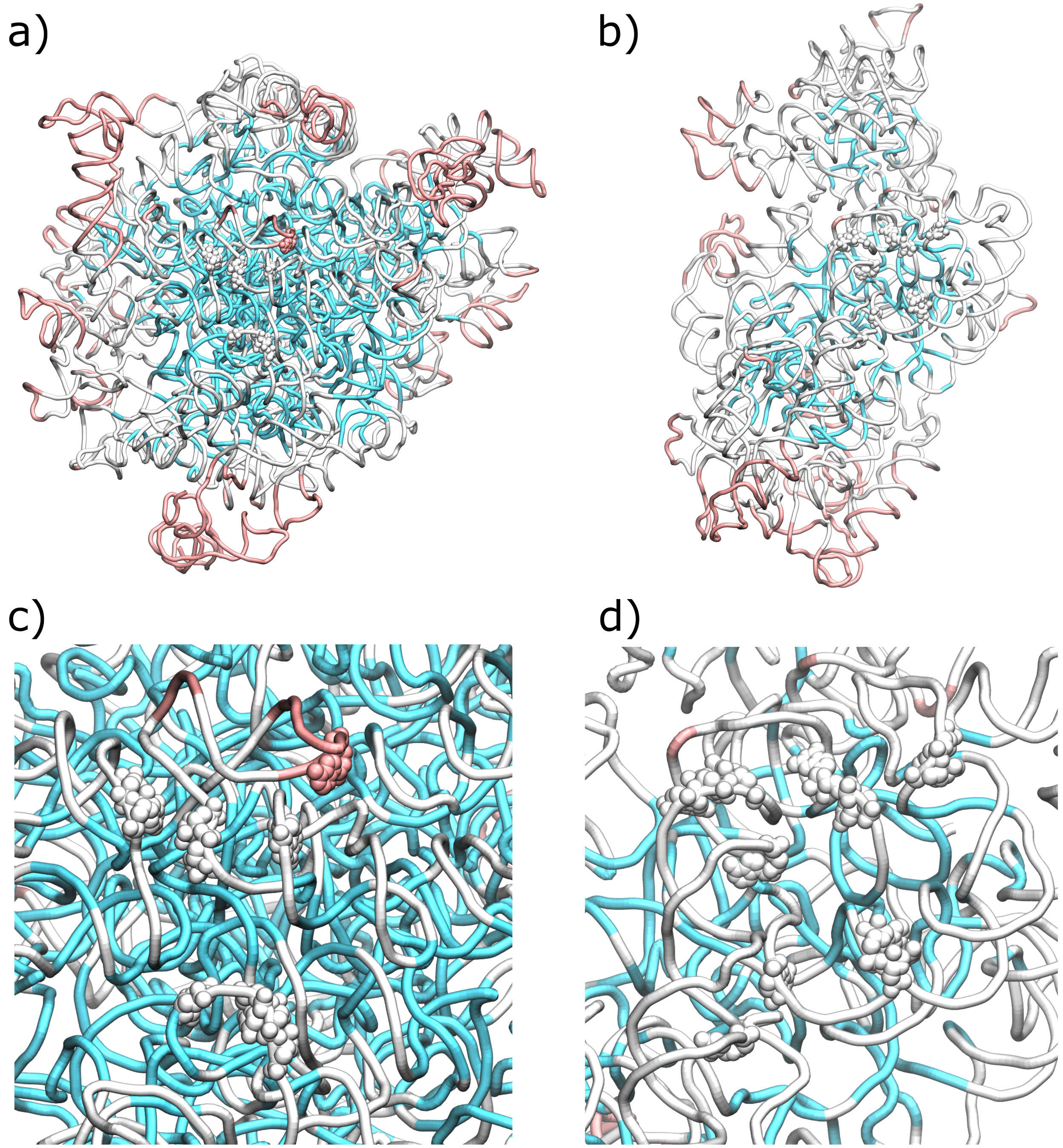


**Figure S9:** Structure of the ribosome, colored by RMSF values, calculated for the unrotated ensemble. a) LSU rRNA, colored blue (rmsf<0.6Å), white (0.6<rmsf<1.2Å) or pink (rmsf >1.2Å). Residues implicated in subunit-bridging ion sites are shown in vdW representation. b) SSU rRNA, colored as in panel a. c) Zoomed-in perspective of subunit-bridging ion sites for LSU rRNA. d) Zoomed-in perspective of subunit-bridging ion sites for SSU rRNA. For both subunits, all residues involved in subunit-bridging ion sites are associated with intermediate (white) or high (pink) flexibility. It is expected that structural methods are unlikely to identify ions in these more dynamic regions, whereas ions have been identified for less mobile (blue) residues[36, 11]. The unrotated ensemble is defined as $(-1.29^\circ < \phi_{\rm body} < 0.12^\circ)$ .

[5] G Torrie and J Valleau. Nonphysical sampling distributions in monte carlo free-energy estimation-umbrella sampling. *Journal of Computational Physics*, Jan 1977.

[30] Y. Marcus. *Ion Properties*. Marcel Dekker, Inc., New York, Basel, 1997.
